# Examining Alzheimer’s Disease modifiable risk factors: Impact of physical activity and diet on neuroanatomy and behaviour in mouse models

**DOI:** 10.1101/2025.02.26.640411

**Authors:** Cindy L. García, Chloe Anastassiadis, Mila Urosevic, Megan Park, Daniel Gallino, Gabriel A. Devenyi, Stephanie Tullo, Yohan Yee, M. Mallar Chakravarty

**Author notes:** Corresponding authors: (CLG), (MMC). These authors contributed equally to this work (Cindy L. García, Chloe Anastassiadis).

## Abstract

Dementia is a global public health challenge, with obesity emerging as an important modifiable risk factor. Here, we examined whether lifestyle interventions can mitigate the effects of diet-induced obesity on body weight, behaviour, and brain anatomy in mouse models. Using a longitudinal design, wild-type and triple-transgenic (3xTgAD) mouse models of Alzheimer’s disease were exposed to a high-fat diet and assigned to one of three interventions: voluntary physical activity, a low-fat diet, and their combination. A high-fat diet significantly increased body weight and induced widespread neuroanatomical changes, with effects modulated by sex and genotype. The combined intervention led to significant weight loss in males of both genotypes. Neuroanatomical analyses revealed that a high-fat diet significantly reduced hippocampal and cerebellar volumes in wild-type mice but had a less pronounced effect on 3xTgAD mice; nevertheless, interventions, particularly the combined approach, increased localized brain volumes in these regions regardless of genotype.

Multivariate integration of behavioural and neuroanatomical measures identified a brain pattern linking hippocampal and cerebellar volumes to intervention and behavioural performance. Spatial gene enrichment analysis of this pattern identified biological processes, including glucose homeostasis, as potential biological mechanisms underlying intervention effects. Overall, these findings suggest that voluntary physical activity and a low-fat diet can modulate brain structure and behaviour, partially counteracting the effects of a high-fat diet, and potentially recruiting biological processes that may support brain health.

## Introduction

Dementia is characterized by severe cognitive decline that impacts the ability to perform tasks related to daily living (Gale et al., 2018), and it is projected to affect ∼78 million people worldwide by 2030 (World Health Organization, 2021). Although increasing life expectancy contributes to the growing prevalence of dementia, age remains the strongest risk factor for neurodegenerative disorders (Niccoli and Partridge, 2012). Importantly, dementia is recognized as a condition that develops over several decades before clinical diagnosis (Iturria-Medina et al., 2016). Consistent with this extended preclinical phase, approximately 15% of dementia cases are attributable to mid-life (45-65 years) modifiable factors such as hypertension, alcohol consumption, and obesity (Livingston et al., 2020).

Strategies aimed at addressing obesity are acknowledged as effective approaches for preventing a range of chronic diseases, such as cancer, diabetes, and cardiovascular conditions (Knowler et al., 1995; Rexrode et al., 1996; Tubiana, 1999). Notably, the Alzheimer’s Association and the Lancet Commission on Dementia report that obesity during mid-life increases the risk of cognitive decline and dementia, whereas this risk is reduced when obesity occurs later in life (Baumgart et al., 2015; Livingston et al., 2024). Several studies have linked weight maintenance and physical activity with a lower risk of developing dementia (Albanese et al., 2017; Hörder et al., 2018; Huang et al., 2022; Zotcheva et al., 2018). For example, combining physical activity with a Mediterranean diet effectively lowers the risk of developing Alzheimer’s Disease (AD), the most common form of dementia (Scarmeas et al., 2009). Conversely, diets high in saturated fats increase inflammatory markers such as interleukin-6 (Milanski et al., 2009), which has been implicated in the pathogenesis of AD (Khan et al., 2020), highlighting the importance of dietary factors in dementia prevention.

Only a few studies have examined the neuroanatomical effects of obesity and intervention strategies. For instance, it has been shown that obesity is associated with lower gray matter volumes in regions including the frontal gyri, insula, and cerebellum (García-García et al., 2019; Gómez-Apo et al., 2021). In previous work from our laboratory, an obesity-inducing high-fat diet led to widespread morphological changes, including a decline in hippocampal volume in the triple transgenic mouse model of Alzheimer’s disease (3xTgAD) (Rollins et al., 2019). In contrast, other studies have found larger volumes in the hippocampus and cerebellum related to physical activity (Brown et al., 2022; Cahill et al., 2015; Raji et al., 2024).

Despite the strong epidemiological evidence linking mid-life obesity to dementia risk, evaluating the effectiveness of intervention strategies in humans remains challenging due to the need for long-term follow-up (Livingston et al., 2024). This emphasizes the importance of experimental models that allow controlled research to understand how risk factors and interventions affect the brain. The present study builds upon our previous work (Rollins et al., 2019) to investigate the effects of three intervention strategies: physical activity, low-fat diet, and the combination of both, in the 3xTgAD mice and their wild-type (WT) counterparts, after receiving a high-fat diet. Animals were studied longitudinally from two to six months of age; a period that corresponds to late adolescence and early adulthood in humans (Cottam et al., 2025). This interval allows for the assessment of intervention efficacy during the initial stages of amyloid and tau development, which is accompanied by cognitive impairments in 3xTgAD mice (Billings et al., 2005).

During this period, mice underwent an interleaved protocol. After being maintained on a standard diet for two months, they were switched to a low-fat or high-fat diet until four months of age, followed by interventions from four to six months. We collected weekly body weight measurements and acquired T1-weighted (T1-w) magnetic resonance images (MRI) of mice at two, four, and six months of age. Behavioural tests (Morris water maze and novel object recognition tests) were conducted after the last MRI scan. We related voxel-wise structural brain changes to behavioural outcomes using partial least squares analysis (PLS), a multivariate method that identifies patterns of covariation between the two datasets without imposing prior neuroanatomical assumptions. We then applied Spatial Gene Enrichment Analysis (SGEA) to gain insight into the potential biological mechanisms underlying the PLS-derived brain pattern. This approach uses spatially resolved gene expression data from the Allen Mouse Brain Atlas (Lein et al., 2007) to identify genes whose expression patterns correlate with the observed brain pattern, and to determine which biological processes are enriched among these genes. The aim of this study was to determine whether lifestyle interventions, including physical activity and a low-fat diet, mitigate the effects of a high-fat diet on body weight, behaviour, and brain anatomy in wild-type and 3xTgAD mice. Our findings support this aim, demonstrating that interventions attenuated high-fat diet effects in a genotype-, sex-, and outcome-dependent manner.

## Results

Raw data and statistical maps generated during this study are available in the repository listed under the “Data and materials availability” section.

### Experimental groups and study timeline

A detailed description of group composition, sex ratios, and diets is provided in Materials and Methods (section ‘Experimental design’). Briefly, all mice were maintained on a standard diet (4% kcal from fat) from birth to 2 months of age (initial N = 165, Fig 1A). From 2-4 months, mice were switched to either a low-fat (10.5% kcal from fat) or a high-fat diet (60% kcal from fat). The low-fat diet is ingredient-matched to the high-fat diet, allowing macronutrient-specific comparisons. WT and 3xTgAD mice that remained on a low-fat diet throughout the experiment are referred to as Controls (WT_Control, Tg_Control). From 4-6 months of age, mice previously exposed to the high-fat diet were assigned to one of four conditions: 1) continuation of the high-fat diet with no further intervention (WT_HighFat, Tg_HighFat); 2) a low-fat diet intervention (WT_Diet, Tg_Diet); 3) access to a wheel for voluntary physical activity while remaining on a high-fat diet (WT_Exercise, Tg_Exercise); or 4) a combined low-fat diet and voluntary exercise intervention (WT_Both, Tg_Both). Body weight was measured weekly from 2 months of age until the end of the experiment. Longitudinal T1-w MRI were acquired at 2, 4, and 6 months of age (100 μm isotropic resolution, Bruker 7T). Following the last scan, novel object recognition and Morris water maze tests were performed to assess intervention effects on memory.

**Fig. 1.**
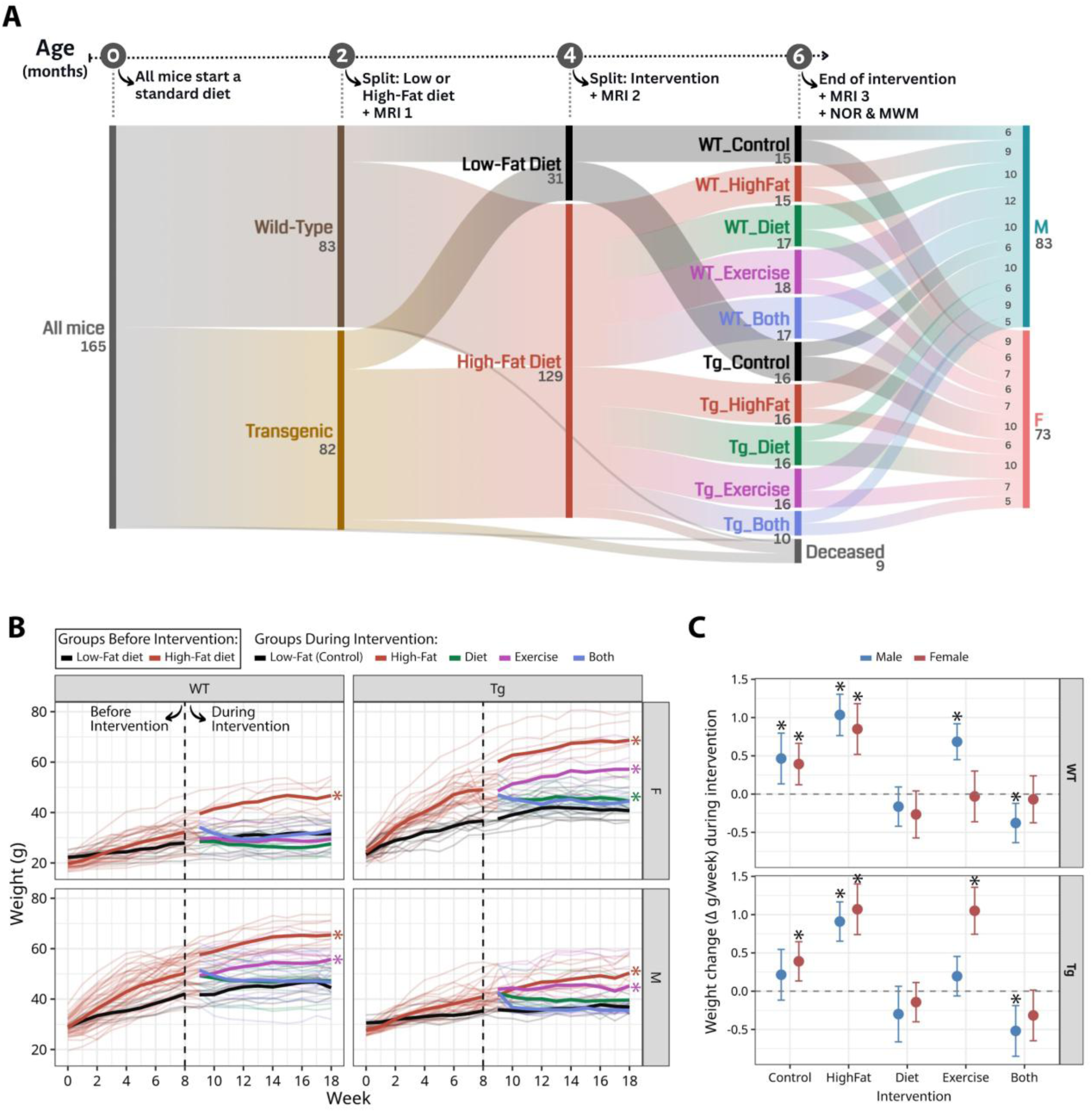
Experimental design and weight changes. (A) For the first two months, all mice received a standard diet. From two to four months of age, mice were put on either a low-fat or a high-fat diet. From four to six months of age, mice were placed in intervention groups. T1w MRIs were collected at two, four, and six months. Behavioural tests (MWM and NOR) were performed after the last MRI scan. (B) Weekly measurements of weight plotted per group. The vertical dotted line indicates the onset of interventions. The x-axis denotes time in weeks, and the y-axis body weight in grams (g). Before interventions, trajectories reflect the weight of mice maintained on a low-fat diet (black) or a high-fat diet (red). After the onset of interventions (week ≥ 8), trajectories are shown separately for each intervention group (see legend). Significant weight differences (p < 0.05) compared against Control are denoted with an asterisk (*). (C) Model-estimated rates of weight change (g/week), stratified by intervention, genotype, and sex. Weight change was estimated using marginal contrasts derived from a four-way linear mixed effects model (*Weight* ∼ *Week*\**Intervention*\**Genotype*\**Sex* + (*Week*|*ID*)). Asterisks (*) indicate estimates with 95% confidence intervals excluding zero (p < 0.05), indicating a significant weight gain or loss. N = 156 mice, with 18 repeated weight measurements each.

### Intervention strategies affect the body weight of mice

Mice were weighed weekly after the change in diet (from standard diet) at two months of age and until the end of the experiment (final N = 156 mice, 18 measures of weight per mouse). To analyze the weight changes per group following intervention onset (Fig 1B), we fit a linear mixed effects model (LME) including a four-way interaction between time (in weeks), intervention, genotype, and sex as fixed effects, along with a random intercept and slope for time at the individual level, to account for repeated measurements and individual variability in weight change over time (*Weight* ∼ *Week*Intervention*\**Genotype*Sex* + (*Week|ID*)). Because higher-order interaction terms are difficult to interpret directly, given that the effect of a variable depends on the values of others (Busenbark et al., 2022), we computed model-based marginal comparisons (Arel-Bundock, 2025). These estimates quantify the expected change in body weight (grams/week) for each intervention, genotype, and sex. This approach enables direct comparison of rates of weight gain or loss across intervention groups (see Statistical Analyses, section ‘Weight and behaviour’). Marginal comparisons (Fig 1C) revealed that during the intervention period (week ≥ 8), mice maintained on a high-fat diet continued to gain weight at a significant rate across genotypes and sexes (0.85 - 1.07 g/week, p < 0.001). Mice in the control groups also kept gaining weight at lower rates than those on a high-fat diet (0.39 - 0.46 g/week, p ≤ 0.006), with 3xTgAD males showing weight stabilization (rates of weight change not significantly different from zero). Interventions differentially altered weight trajectories depending on sex and genotype. Mice receiving a Diet intervention showed weight stabilization. Exercise alone resulted in varied outcomes. WT males (0.68 g/week, 95% CI [0.45, 0.92], p < 0.001) and 3xTgAD females (1.05 g/week, 95% CI [0.74, 1.36], p < 0.001) continued to gain weight despite access to a wheel while maintained on a high-fat diet. In contrast, WT females and 3xTgAD males displayed weight stabilization. The combined intervention (Both) resulted in significant weight loss in males of both genotypes (WT: -0.38 g/week, 95% CI [-0.63, -0.121], p = 0.004; 3xTgAD: -0.52 g/week, 95% CI [-0.85, -0.19], p = 0.002), while females of both genotypes exhibited weight stabilization. Taken together, these results point to a sex- and genotype-dependent modulation of weight trajectories in response to intervention.

### Volumetric effects of genotype, high-fat diet, and interventions

To assess the effects of genotype, high-fat diet, and interventions on brain anatomy, we applied a hierarchical LME model to the 182 regional volumes derived from the MAGeT brain segmentation algorithm (Chakravarty et al., 2013) using the Dorr-Steadman-Ullmann-Richards-Qiu-Egan (DSURQE) mouse brain atlas (Dorr et al., 2008; Richards et al., 2011; Steadman et al., 2014; Ullmann et al., 2013), as described in Materials and Methods, section ‘Volumetric analysis’. We used brain images collected at 2, 4, and 6 months of age. The model included an interaction between group and age, modeled as a quadratic term to account for nonlinear trajectories (Fjell et al., 2010), sex as an independent variable, and a random intercept per mouse (model: *Volume* ∼ *Group*\*poly(*Age*,2) + *Sex* + (*1|ID*)). All model outputs were corrected for multiple comparisons at a 10% False Discovery Rate (FDR) threshold (Benjamini and Yekutieli, 2001).

Comparing 3xTgAD and WT mice under control conditions revealed widespread volumetric differences, as detailed in the Supplementary Results (section ‘Volumetric analysis’). However, when examining the impact of the high-fat diet and subsequent interventions, significant volumetric changes were restricted to WT groups. In the 3xTgAD mice, these regional volumetric measures showed limited detectable changes (Supplementary Figs S1-S3).

These findings suggest that while regional volumetric analysis is highly effective for capturing global genotype differences, it may be less sensitive to more subtle changes, such as early atrophy or partial recovery in 3xTgAD mice. This motivated our subsequent use of Deformation-Based Morphometry (DBM), a voxel-wise approach capable of identifying localized structural shifts that may be obscured when averaging across entire anatomical regions (Germann et al., 2025; van Eede et al., 2013).

### Deformation-based morphometry identifies similar patterns of local brain changes

To capture changes without relying on predefined neuroanatomical boundaries, we performed voxel-wise DBM analysis (see Materials and Methods, section ‘Deformation-Based Morphometry analysis’). DBM quantifies local structural differences by measuring the deformations required to align individual brain images to a common population template. These voxel-wise deformations are summarized by the Jacobian determinant, which reflects local volume expansion (positive values) or contraction (negative values) relative to the population average (Chung et al., 2001). DBM has been previously applied by our group to study morphological changes in the 3xTgAD (Kong et al., 2018), 3xTgAD receiving a high-fat diet (Rollins et al., 2019), models of synucleinopathy (Tullo et al., 2024), and neurodegeneration in Parkinson’s and frontotemporal dementia (Bussy et al., 2023; Tullo et al., 2025).

LME models were fitted at each voxel using the log-transformed Jacobian determinant as outcome, with an interaction between group and age in days (modeled as a quadratic polynomial), and sex, along with a random intercept for each mouse (model: *log-Jacobian* ∼ *Group*\*poly(*Age*,2) + *Sex* + (*1|ID*)). Data from all time points were included, and results were corrected for multiple comparisons with FDR at a 10% threshold. While the quadratic interaction term was included to detect potential non-linear trajectories, these effects were limited to highly localized cortical areas and the olfactory bulbs (Supplementary Fig S4). In contrast, the linear term of the interaction captured widespread changes in brain volume over time (Fig 2). These linear changes are most relevant to our hypothesis, which anticipated gradual neuroanatomical changes driven by a high-fat diet and the subsequent interventions.

**Fig. 2.**
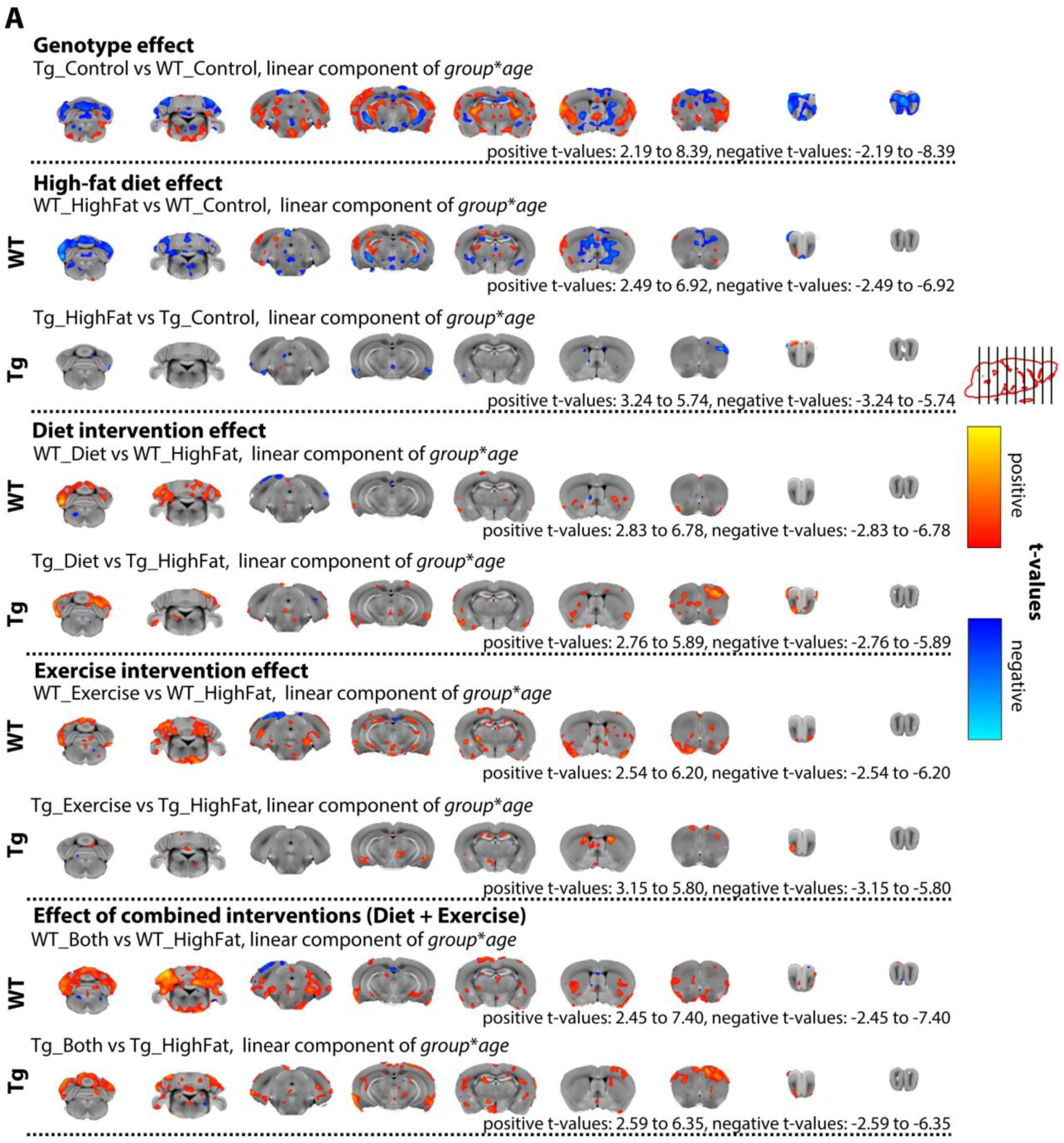
Age-dependent group differences in brain volume change. The *t*-statistic maps show the linear component of the group and age interaction, thresholded at 10% FDR and displayed at their peak *t*-values, overlaid on the population-average brain. Estimates were derived from a linear mixed-effects model of the log-transformed Jacobian determinant (∼ *Group* * poly(*Age*, 2) + *Sex* + (1 | ID)). N = 152 mice.

DBM results are summarized in Fig 2. The comparison between Tg_Control versus WT_Control (genotype effect) revealed widespread significant differences in the cerebellum, hippocampus, caudoputamen, cortex, and olfactory bulbs. High-fat diet effects (comparison of Tg_HighFat and WT_HighFat against their controls) were again found in the cerebellum as well as the hippocampus and caudoputamen, with no major voxel-wise effects detected in 3xTgAD mice, consistent with the volumetric analysis. This suggests that the high-fat diet effect is modulated by genotype, with WT mice exhibiting greater sensitivity.

However, while the effects of a high-fat diet are less prominent in the 3xTgAD mice, the subsequent voxel-wise analysis of intervention effects revealed localized structural growth that was previously obscured in the regional volumetric analysis.

Relative to mice fed a high-fat diet, the dietary intervention (Fig 2, ‘Diet intervention effect’) was associated with a significant volume increase within cerebellar regions in both genotypes, reaching control-like values, whereas physical activity (Fig 2, ‘Exercise intervention effect’) promoted volume growth in hippocampus and cerebellum. The combined intervention (Fig 2, ‘Effect of combined intervention effect’) resulted in significant structural growth in the cerebellum of both genotypes and in the hippocampus of 3xTgAD mice. No significant sex differences were observed across any comparisons. Evidence regarding the directionality of these structural changes is provided in the following section, which illustrates the significant peak voxel trajectories (Figs 3-5 and Supplementary Figs S5-S6).

**Fig. 3.**
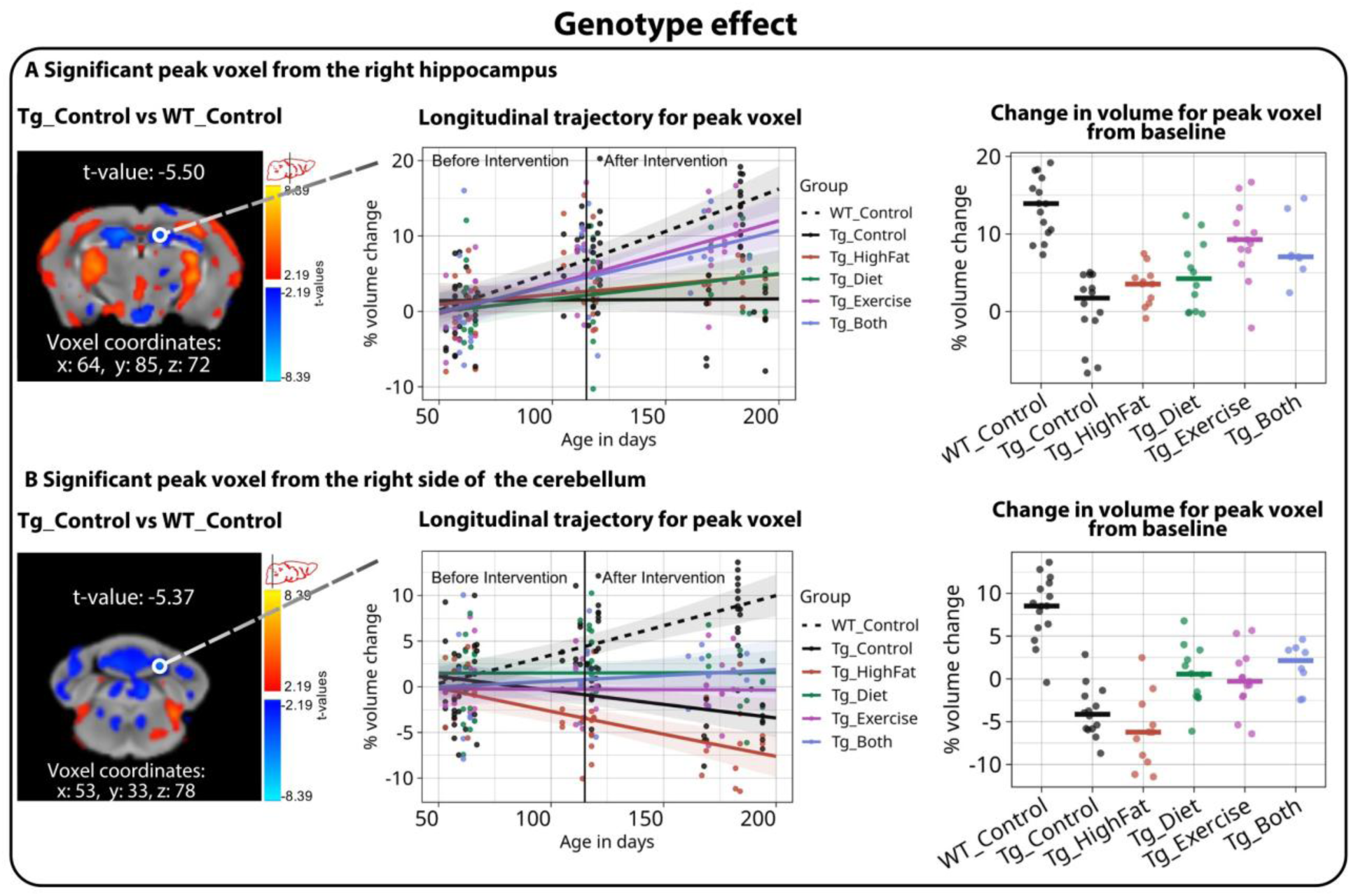
Trajectories of significant peak voxels identified for the genotype effect. The figure shows peak voxels selected from the comparison between Tg_Control and WT_Control, which highlights the genotype effect on the hippocampus (A) and cerebellum (B). The left panels show the statistical maps of the interaction between the group and age in days, thresholded at 10% FDR, together with the corresponding *t*-values and voxel coordinates (from model: *log-Jacobian* ∼ *Group*\*poly(*Age*,2) + *Sex* + (*1|ID*)). The middle panels show the percent change in volume from baseline in the selected peak voxels. The right panels show the percent volume change from baseline at the final time point; horizontal lines represent the group median, and individual data points are overlaid to illustrate distribution. FDR: False Discovery Rate.

In summary, DBM analysis demonstrates that genotype-driven differences follow a consistent spatial signature involving the cortex, cerebellum, and hippocampus, a pattern that is partially mirrored in WT mice exposed to a high-fat diet. Importantly, by bypassing the constraints of predefined anatomical boundaries, DBM successfully captured intervention-related structural preservation in 3xTgAD mice that were not detectable using regional volumetry.

### Peak-voxel trajectories reveal that intervention strategies influence local brain structure

Thus far, both volumetric and DBM analyses revealed statistical effects driven by genotype, high-fat diet, and intervention strategies. To characterize the directionality of the volumetric changes underlying these effects, we selected peak voxels to examine voxel-wise trajectories. Peak voxels were defined as those with the highest t-values from the DBM statistical maps that survived a 10% FDR threshold.

In Fig. 3, the left panels display the statistical maps overlaid on the population average, along with the t-value and coordinates of the selected peak voxel. The middle panels show longitudinal trajectories expressed as percent volume change from baseline. The right panels display the percent volume change reached at the end of the intervention period (6 months of age), highlighting the group differences observed at the final time point.

We first illustrate the genotype effect by comparing Tg_Control and WT_Control at significant peak voxels identified in two brain regions: the hippocampus (Fig. 3A) and the cerebellum (Fig. 3B). In the hippocampus, by the final time point (right panels), Tg_Control shows minimal volumetric change from baseline (∼2%), which is significantly lower than the ∼20% increase observed in WT_Control. In the cerebellum, Tg_Control exhibits a significant reduction in volume (∼ -4%), whereas WT_Control shows an average increase of ∼8% at the same voxel. These patterns are consistent with the volumetric results (Supplementary Figs. S1B–S1C). Trajectories for the same voxel are also shown for the other intervention groups for visual comparison.

To examine the effect of a high-fat diet, we identified significant peak voxels showing differences between HighFat and Control groups. In WT mice, a peak voxel in the hippocampus (Fig. 4A, top panel) shows that WT_Control mice exhibit ∼20% volume growth from baseline by the final time point, whereas WT_HighFat mice remain close to baseline, indicating markedly reduced growth. In the cerebellum (Fig. 4B, top panel), WT_HighFat mice display an ∼7% reduction in volume relative to baseline, contrasting with the modest volume increase observed in WT_Control mice. In 3xTgAD mice, a significant peak voxel in the cerebellum (Fig. 4A, bottom panel) reveals divergent trajectories between Tg_Control and Tg_HighFat groups, with Tg_HighFat mice showing a pronounced decline in volume across time. Together, these results indicate that a high-fat diet alters longitudinal volume trajectories, with particularly pronounced effects observed in WT mice.

**Fig. 4.**
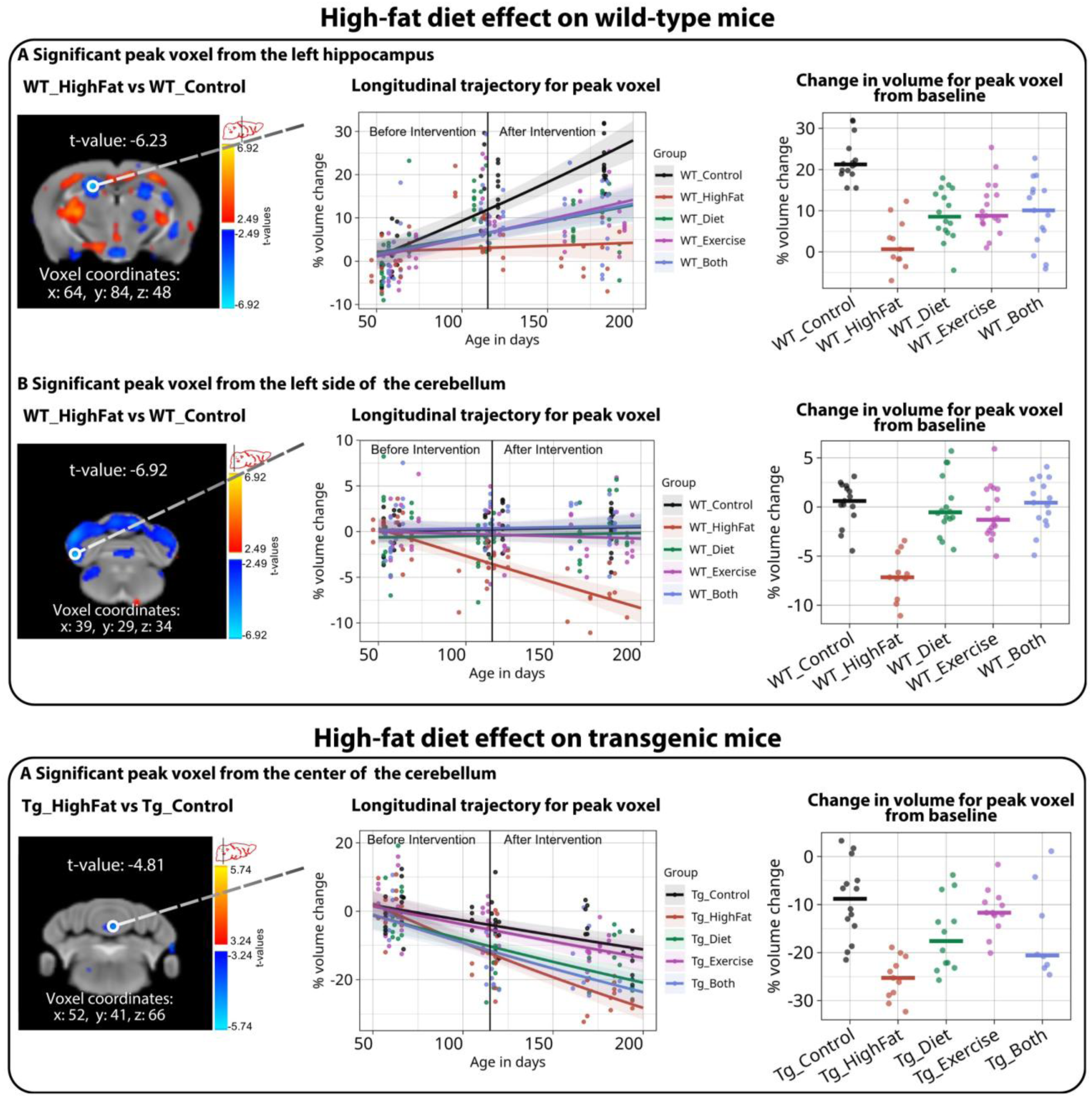
Trajectories of significant peak voxels identified for the high-fat diet effect. Significant peak voxels were selected from comparisons between WT_HighFat and WT_Control (top panels, A-B) and between Tg_HighFat and Tg_Control (bottom panel, A), highlighting genotype-specific effects of the obesity-inducing high-fat diet. Left panels: statistical maps of the interaction between group and age (from model: *log-Jacobian* ∼ *Group*\*poly(*Age*,2) + *Sex* + (*1|ID*), 10% FDR corrected). Middle panels: longitudinal percent change in volume from baseline across all three time points. Right panels: percent volume change from baseline and median at the last time point. FDR: False Discovery Rate.

Lastly, we report peak voxels identified for each genotype under the combined intervention (Fig. 5). In WT mice, a significant peak voxel in the cerebellum (Fig. 5A, top panel) shows that WT_Both mice exhibit greater percent volume change from baseline (∼10%) than WT_Control and differ significantly from WT_HighFat, which shows a decline in volume. This pattern is consistent with enhanced regional volume growth under the combined exercise and diet intervention.

**Fig. 5.**
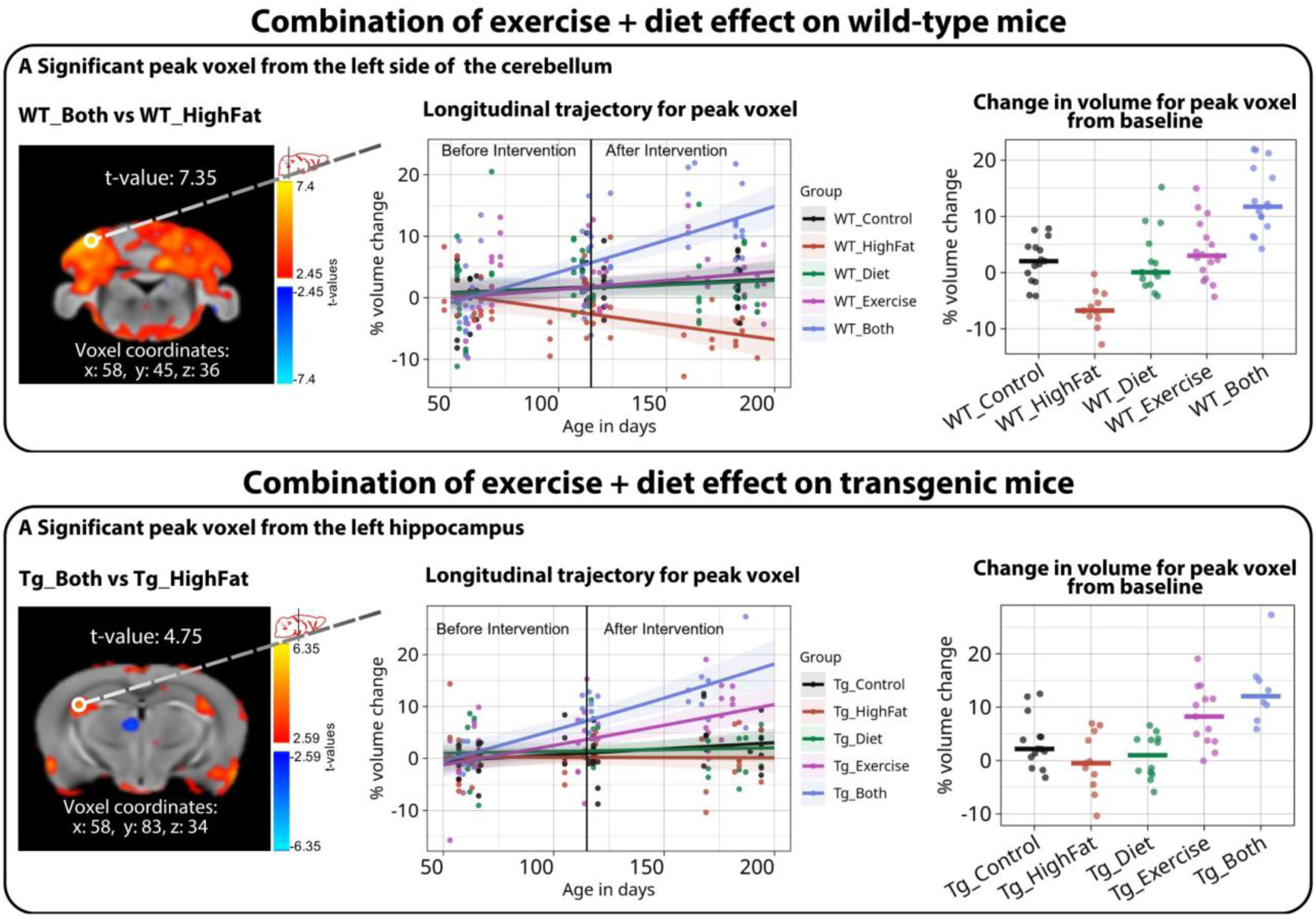
Trajectories of significant peak voxels identified for the combined interventions. Significant peak voxels were selected from the left side of the cerebellum, from the comparison between WT_Both and WT_HighFat (top panel, A), and the left hippocampus from the comparison between Tg_Both and Tg_HighFat (bottom panel, A), highlighting effects of the combined exercise and diet intervention in each genotype. Panels show the statistical maps of the interaction between group and age (from model: *log-Jacobian* ∼ *Group*\*poly(*Age*,2) + *Sex* + (*1|ID*), 10% FDR corrected, left panels), longitudinal percent change in volume from baseline across all three time points (middle panels), and percent volume change from baseline and the group median at the last time point (right panels). FDR: False Discovery Rate.

In 3xTgAD mice, a peak voxel in the hippocampus (Fig. 5A, bottom panel) reveals a similar pattern: Tg_Both mice show greater percent volume change from baseline (∼10%) than Tg_Control, indicating localized volume growth under the combined intervention despite the absence of a detectable high-fat diet effect in this region.

Additional peak voxel trajectories for diet and physical activity groups are shown in Supplementary Figs. S5-S6, where similar trends are observed, with intervention strategies generally promoting volume growth compared with mice that did not receive an intervention.

### Intervention effects on memory performance

After the final MRI scan, mice underwent the novel object recognition (NOR) and Morris water maze (MWM) tests (see Materials and Methods, ‘Behavioural tests’) to assess recognition and spatial memory differences between groups. In the NOR test, mice were exposed to familiar and novel objects. Because mice prefer novelty, if the mouse remembers the familiar object, it will spend more time exploring the novel object (Lueptow, 2017). This paradigm assesses short-term recognition memory, which relies on the perirhinal cortex and dorsal hippocampus (Antunes and Biala, 2012). In contrast, the MWM assesses long-term spatial memory dependent on the hippocampus (Bromley-Brits et al., 2011). In the test, mice are trained to find a hidden escape platform in a water-filled tank and are expected to remember the platform’s location based on spatial cues in the room during the probe trial (Bromley-Brits et al., 2011). Variables of interest for the NOR included the percentage of time spent near the novel object and the discrimination index (Antunes and Biala, 2012), which quantifies preference for the novel object. It is calculated as the difference in time spent exploring the boundary of the novel versus familiar objects, divided by total exploration time, with higher values indicating better recognition memory. For the MWM, Gallagher’s cumulative measure (GCM) (Pereira and Burwell, 2015) and the number of pass-throughs were analyzed. Lower GCM values reflect more efficient learning, while higher counts of passages near the location where the platform was previously located (pass-throughs) indicate better spatial memory (Maei et al., 2009). These outcomes represent standard measures commonly reported in the literature. Under standard chow conditions, MWM deficits typically emerge around 6 months (Billings et al., 2005; Webster et al., 2014), with NOR impairments appearing near 9 months (Clinton et al., 2007; Webster et al., 2014). Although exposure to a high-fat diet can shift and accelerate this trajectory (Kim et al., 2017; Knight et al., 2014; Sah et al., 2017), reported outcomes remain inconsistent due to variations in feeding duration and experimental windows.

Behavioural outcomes were analyzed using regression models appropriate for each variable, with sex and genotype included as modeled factors. For each outcome, competing models with and without higher-order interactions were compared to determine the most appropriate representation of the data (for NOR: *Behavioural Outcome* ∼ *Intervention* * *Genotype* * *Sex* or *Behavioural Outcome* ∼ *Intervention* * *Genotype* + *Sex*. For MWM: *Behavioural Outcome* ∼ *Intervention* * *Genotype* * *Sex + Software + Weight* or *Behavioural Outcome* ∼ *Intervention* * *Genotype* + *Sex + Software + Weight*). For details, see Statistical Analyses, section ‘Weight and behaviour’. As a result, intervention effects are reported as model-based marginal contrasts, which estimate the direction and magnitude of expected change in each behavioural outcome under a given intervention, relative to Control, while adjusting for the covariates in the selected model. Confidence intervals that include zero indicate that the estimated intervention effect is not statistically different from the Control.

In the NOR test, WT males in the Exercise intervention showed reduced exploration time (p = 0.02) and increased immobility (p = 0.01), whereas WT females receiving the combined intervention travelled less (p = 0.009) and spent more time immobile (p = 0.02). These patterns suggest altered activity or anxiety-like behavior rather than global memory deficits (Supplementary Table S1).

In the MWM, a high-fat diet alone did not impair spatial memory at six months, as reflected by GCM and pass-throughs (Fig. 6A-B). In contrast, interventions improved performance in several sex- and genotype-specific combinations. WT males in the combined intervention and WT females in the Diet intervention showed reduced GCM (p < 0.05), indicating more efficient learning. Males from both genotypes in the combined intervention also exhibited increased pass-throughs (p < 0.01), consistent with improved spatial memory. Female mice displayed more selective effects, including increased pass-throughs in WT Exercise (p = 0.04) and reduced performance in 3xTgAD females under the Diet intervention (p = 0.003) (Fig. 6A-B).

**Fig. 6.**
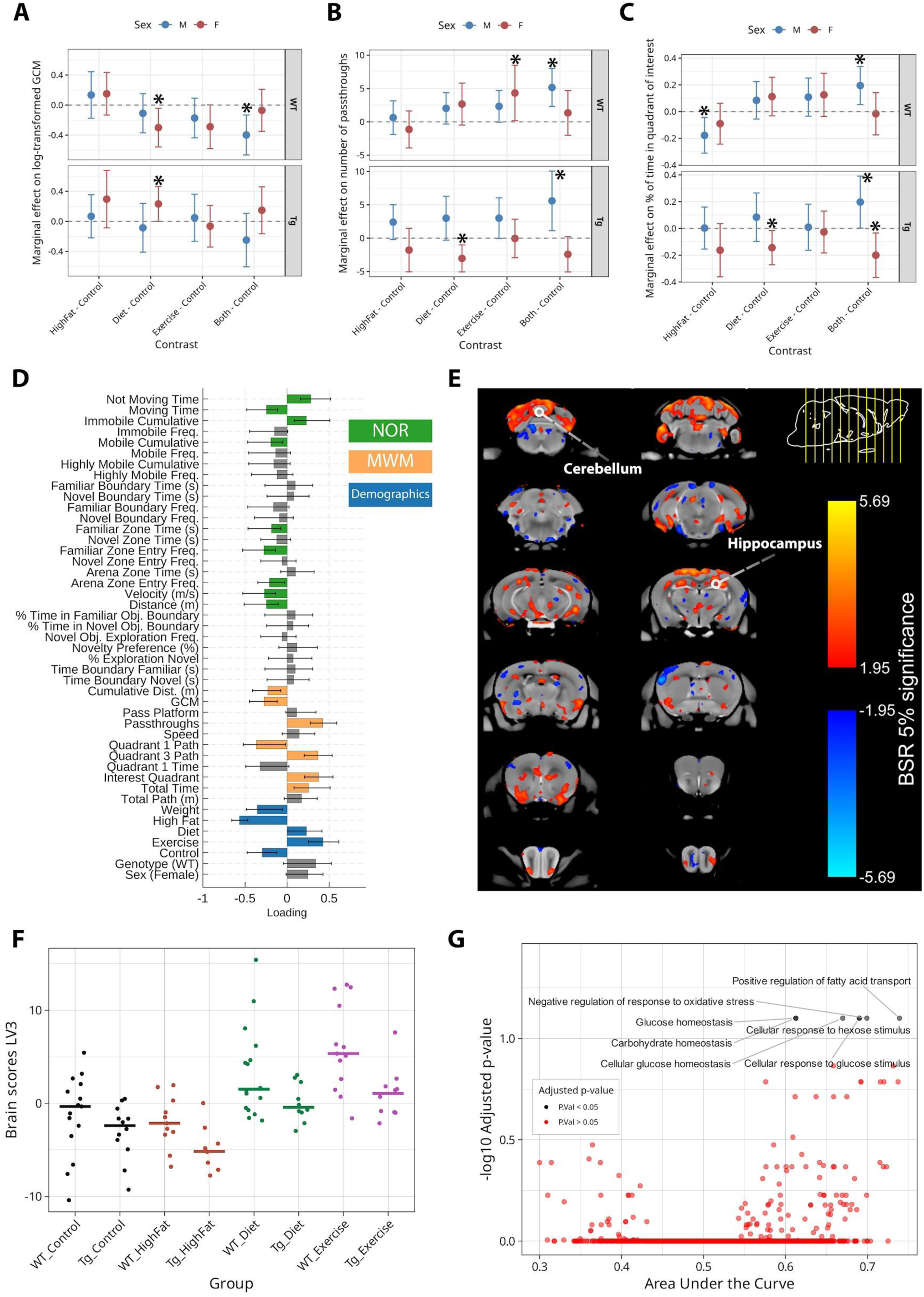
Intervention-associated effects across behavioural outcomes, multivariate, and gene enrichment analyses. (A) Marginal contrasts for log-transformed Gallagher’s Cumulative Measurement (GCM), where lower GCM values indicate better learning. (B) Marginal contrasts for the number of pass-throughs, a measure of spatial memory retention; higher values indicate better performance. (C) Marginal contrasts for the percentage of time spent in the target quadrant of interest (where the platform was located during training); higher values indicate better performance. Points show contrasts comparing each intervention (HighFat, Diet, Exercise, and Both) to the Control, stratified by Genotype (WT, top; Tg, bottom) and Sex (Blue = Male, Red = Female). Error bars denote 95% confidence intervals; intervals crossing the zero line indicate non-significant differences. Tg = 3xTgAD mice. (D) shows the PLS-LV3 behavioural pattern that maximally covaries with the brain pattern (E). For the behavioural data (D), variables from the NOR (in green), the MWM (in yellow), and the treatment groups or demographics (in blue) are shown. The bars show how much each behaviour contributes to the pattern of LV3 (x-axis: loading values). Singular value decomposition estimated the size of the bars, whereas confidence intervals were estimated by bootstrapping. Bars with error bars that crossed the zero line are not significant (in gray). (E) Brain loading BSRs are overlaid on the population average. BSR thresholded to 95% confidence interval (N = 96 mice with complete data). (F) Group-wise distribution of brain scores (derived from PLS) and median, illustrates that individuals in the Diet and Exercise intervention are highly represented by the LV3 pattern. (G) Scatter plot showing significant modules identified with SGEA. Seven gene ontology (GO) terms were associated with top-ranked genes, defined as those most strongly correlated with the LV3 brain loadings, with an area under the curve > 0.5 and P < 0.05. The significant terms are: negative regulation of response to oxidative stress (GO:1902883), glucose homeostasis (GO:0042593), carbohydrate homeostasis (GO:0033500), cellular glucose homeostasis (GO:0001678), cellular response to hexose stimulus (GO:0071331), cellular response to glucose stimulus (GO:0071333), and positive regulation of fatty acid transport (GO:2000193). PLS: Partial Least Squares. SGEA: Spatial Gene Enrichment Analysis.

Quadrant occupancy further highlighted sex- and genotype-specific effects (Fig. 6C). WT males in the combined intervention spent 19.5% more time in the target quadrant (the location of the platform during training), indicating improved performance (p = 0.007), whereas WT males on a high-fat diet spent 17.7% less time in this quadrant (p = 0.009), reflecting poorer performance. In 3xTgAD females, the combined intervention and Diet reduced time in the target quadrant by 20.0% and 14.0%, respectively (p < 0.05), while 3xTgAD males in the combined intervention showed a 19.0% increase (improved performance) (p = 0.047).

Together, these findings suggest that a high-fat diet did not produce widespread memory deficits at this age, but that interventions can modulate memory-related outcomes in a sex- and genotype-dependent manner, with males generally benefiting from the combined intervention.

### Multivariate analyses of neuroanatomical changes, interventions, and behaviour

Given the high dimensionality of our data, which include 10 groups, thousands of voxels, and over 30 behavioural variables, we used Partial Least Squares (PLS) analysis, a multivariate method that identifies relationships explaining shared variance across datasets (Rännar et al., 1994). In this study, we applied behavioural PLS, in which brain measurements (voxel-wise log-Jacobian values) and behavioural variables (derived from MWM and NOR tests, intervention assignment, and demographics) were used as inputs. This approach enables dimensionality reduction while preserving the covariance structure of the data, allowing detection of patterns that may not be apparent when variables are examined individually. Mice in the combined intervention groups were excluded due to missing behavioural data (see Materials and Methods, ‘Partial Least Squares analysis’).

PLS treats each of the two input matrices, here behaviour and brain measurements, as “blocks” and iteratively transforms them to find the maximum overlap between the two (maximized shared variance) (Gerlach et al., 1979). For details on these transformations, refer to McIntosh and Misic, 2013 (McIntosh and Mišić, 2013). PLS produces several latent variables (LV), each of which encompasses a combination of one brain pattern and one behavioural pattern that maximally covary together and that explain a percentage of the covariance among the data. The LVs are ordered according to the amount of covariance they explain, such that the first LV (LV1) captures the largest overall shared variance between behaviour and brain measures. However, subsequent LVs capture additional orthogonal patterns of covariation independent of earlier LVs and may reflect relationships, including intervention-related effects, that are not dominant at the global level. To determine whether each LV represented a statistically significant brain-behaviour relationship rather than a chance association, significance was assessed using permutation testing (3000 permutations), while standard errors and confidence intervals were estimated via bootstrap resampling (3000 samples). Voxel-wise bootstrap ratios (BSRs) were then used to identify significant brain regions contributing to each LV. BSRs were thresholded at ±1.96, corresponding to a 95% confidence interval (see Statistical Analyses, section ‘Partial Least Squares analysis’). While permutation-based inference is a standard approach, recent work from our laboratory (Danyluik et al., 2024) recommends that the significance of LVs be evaluated using measures beyond permutation-based p-values. In line with those findings and field standards (McIntosh and Lobaugh, 2004), we interpreted the LVs based on the convergence of three factors: statistical significance, the amount of covariance explained (LV strength), and the reliability of its spatial patterns across bootstrap resampling. This approach ensures that our reported results reflect robust patterns rather than artifacts of the permutation procedure.

A total of 45 latent variables (LVs) were obtained, corresponding to the number of behavioural measures analyzed. Permutation testing identified four significant LVs (LV1–LV4; p = 0.0170, 0.0010, 0.0010, and 0.0013), which explained 29.36%, 13.44%, 12.46%, and 7.68% of the total covariance, respectively. The significance and covariance explained by all 45 LVs are shown in Supplementary Fig S8.

LV1 explained 29.36% of the covariance (Supplementary Fig S9A). It was characterized by positive brain loadings (positive BSR) in the cerebellum, olfactory bulbs, and hippocampus. This brain pattern was associated with wild-type mice, lower body weight, and better memory performance, representing the healthiest mice with no AD-like characteristics.

LV2 accounted for 13.44% of the covariance (Supplementary Fig S9B) and displayed negative loadings in the cerebellum and hippocampus, alongside widespread increases in the caudoputamen and cortex. This pattern correlated with male 3xTgAD mice, higher body weight, and poorer memory, reflecting mice at risk of developing AD-related features.

LV3 explained 12.46% of the covariance (Fig 6D-E) and was characterized by positive BSR in the hippocampus and cerebellum. It correlated positively with low-fat diet and physical activity interventions, negatively with no intervention and higher weight, and with better cognitive performance on the MWM (GCM, pass-through, and time in the target quadrant). LV3 captured intervention-related effects in a data-driven, anatomically unconstrained way, highlighting widespread changes including the hippocampus and cerebellum, and aligned with patterns observed in volumetric and voxel-wise analyses.

LV4 explained 7.68% of the covariance (Supplementary Fig S9C) and displayed negative and positive BSRs across multiple regions, including positive loadings in the olfactory bulbs. This LV was associated with wild-type mice.

Given that LV3 was the only LV capturing a correlation between brain volume changes and intervention strategies, we examined brain scores, which reflect how strongly each mouse expresses the brain pattern of an LV. Corresponding behaviour scores indicate how well each mouse exhibits the behavioural pattern of the LV. Brain scores for LV3 were plotted against body weight, GCM, and number of pass-throughs.

A negative correlation with weight (r = -0.35; Supplementary Fig S10) indicated that heavier mice were less likely to exhibit the LV3 brain pattern, suggesting an inverse relationship between cerebellar and hippocampal volumes and body weight. Brain scores also negatively correlated with GCM (r = -0.27; Supplementary Fig S11A), consistent with better spatial memory performance, and positively correlated with pass-through counts (r = 0.42; Supplementary Fig S11B), indicating improved learning and memory of the platform’s location.

Lastly, we plotted brain scores per group (Fig 6F) and found higher brain scores for groups that received diet and exercise interventions, meaning the brain pattern of LV3 was better representing mice in the physical activity and low-fat diet groups. Plots of group-wise brain scores for other LVs support that LV1 is better at representing WT mice (Supplementary Fig S12A), LV2 is better at representing 3xTgAD mice, with a higher brain score median for 3xTgAD mice that didn’t receive any intervention (Tg_HighFat, Supplementary Fig S12B), and LV4 is capturing features of WT mice (Supplementary Fig S12C).

### Biological processes underlying the anatomical signature of LV3

Given that the LV3 brain pattern was associated with intervention strategies, we hypothesized that these interventions contributed to the observed structural pattern. This hypothesis is grounded in prior evidence that these interventions influence neuroanatomy (Brown et al., 2022; Jensen et al., 2021; Raji et al., 2024) and metabolic pathways (Gondim et al., 2015; Macedo et al., 2020).

Under this assumption, we examined the biological processes potentially driving LV3 by comparing its spatial brain pattern with gene expression data from the Allen Mouse Brain Atlas (Arnatkevičiūtė et al., 2019; Lein et al., 2007; Wang et al., 2020). Genes were ranked by Spearman correlation, followed by a spatial gene enrichment analysis (SGEA) to statistically identify overrepresented biological processes (GO terms) (Reijnders and Waterhouse, 2021), whose gene expression patterns resemble the LV3 spatial brain pattern. See Materials and Methods, section ‘Spatial Gene Enrichment Analysis’ for details.

Seven significant GO terms emerged (Fig 6G), highlighting key biological processes: 1) Negative regulation of response to oxidative stress, a process that reduces, prevents, or stops responses to oxidative stress (GO:1902883); 2) Glucose homeostasis, which refers to the maintenance of a stable internal glucose level within an organism or cell (GO:0042593); 3) Carbohydrate homeostasis: the regulation of stable carbohydrate levels within an organism (GO:0033500); 4) Cellular glucose homeostasis: the maintenance of glucose levels specifically within a cell (GO:0001678); 5-6) Cellular response to hexose/glucose stimulus: a process where cells alter their state or activity in response to hexose/glucose (GO:0071331 and 0071333); and 7) Positive regulation of fatty acid transport: a process that increases or activates the transport of fatty acids (GO:2000193).

The ancestry analysis of cellular response to glucose stimulus (GO:0071333) (EMBL-EBI, n.d.) revealed hierarchical relationships with carbohydrate homeostasis, glucose homeostasis, cellular glucose homeostasis, and cellular response to hexose stimulus. This suggests a coordinated regulatory network controlling glucose metabolism. In contrast, negative regulation of oxidative stress response and positive regulation of fatty acid transport were independent, indicating distinct pathways.

## Discussion

While there are no disease-modifying treatments for AD, reducing the incidence of new cases through intervention strategies emerges as an important approach to addressing this global challenge (Alwan et al., 2024). In this manuscript, we leveraged neuroimaging techniques to study the effects of a high-fat diet on the mouse brain and how intervention strategies, including physical activity, a low-fat diet, or their combination, can mitigate these effects. We collected a unique dataset encompassing 156 mice, divided into similar groups per genotype (3xTgAD and WT), balanced for sex. In these groups, we evaluated weight, behaviour, and neuroanatomy, and used multivariate analysis to find hidden patterns in the data that led to the discovery of possible biological processes behind the impact of exercise and a change to a low-fat diet. Although our focus was not to understand how interventions delay or prevent dementia, our findings support their role in promoting brain health by reversing the effects of a high-fat diet in mice.

To evaluate neuroanatomical changes, we used volumetric and DBM analyses. DBM provides insights into subtle, region-specific deformations that volumetric measures may overlook. We found that genotype itself affected the cortex volume, as found before in previous work done in our lab (Kong et al., 2018; Rollins et al., 2019). In addition, we found pronounced changes in the olfactory bulbs, hippocampus, caudoputamen, and cerebellum, which replicate work from our lab aimed at characterizing the neuroanatomical trajectories of the 3xTgAD mice (Kong et al., 2018). We also found that a high-fat diet affected the hippocampus and cerebellum (in WT mice only), structures that have been shown to present modifications with a high-fat diet or obesity in both rodents and humans, impacting neurogenesis and synaptic plasticity in the hippocampus in rodents (González Olmo et al., 2023; Lindqvist et al., 2006), and causing decreased volume in the left cerebellum in both adults and children (Herrmann et al., 2019; Ou et al., 2015). The lack of effects of a high-fat diet on 3xTgAD mice could be attributed to their genetic predisposition, which is linked to neuroanatomical vulnerabilities (Esquerda-Canals et al., 2013; Gama Sosa et al., 2010; Kong et al., 2018). This genetic vulnerability could explain why we found smaller volumes for the cerebellum and hippocampus in 3xTgAD mice.

Peak voxels allowed us to understand the effect of intervention strategies. In many instances, we noticed that the interventions affected the voxel-wise volumes (either expansions or contractions) to comparable levels as the control groups, even exceeding those levels for the groups that received a combined intervention. In animal models, a high-fat diet increases microglial activation (Knight et al., 2014), causes changes in hippocampal mitochondrial morphology, and leads to a reduction in synapse number (Martins et al., 2017). On the contrary, voluntary physical activity has been shown to increase dendritic complexity in the dentate gyrus (Redila and Christie, 2006), regulate synaptic transmission in the hippocampus (Vaynman et al., 2006), promote angiogenesis after an ischemic injury (Tang et al., 2018), and increase the volume of both hippocampus and cerebellar regions (Cahill et al., 2015), while a low-fat diet has not been well studied in terms of effects at the microscopic level. Of interest, a study evaluating the same interventions as in this work found that they mitigate high-fat diet-related amyloid-β accumulation and memory impairments in transgenic mice that overexpress the human amyloid-β precursor protein (APP) in the hippocampus (Maesako et al., 2012).

We studied the 3xTgAD mice to model AD-relevant susceptibility to risk factors. These mice have transgenes for APPSWE and PS1M146V, along with an extra mutation associated with frontotemporal dementia, tauP301L, resulting in the formation of both Aβ plaques and neurofibrillary tangles in the model (Oddo et al., 2003). Previous studies indicate that, by 6 months of age, these mice exhibit early hippocampal pathology, with some amyloid-β accumulation and tau phosphorylation (Belfiore et al., 2019; Jullienne et al., 2022). Importantly, in line with prior literature, spatial memory deficits assessed by the MWM can appear as early as 6 months, whereas recognition memory impairments assessed by the NOR test tend to emerge later, around 9 months (Webster et al., 2014). Consistent with these expectations, we observed behavioural effects in the MWM at 6 months, whereas NOR performance remained largely unaffected. Notably, these effects were selective for genotype and sex. These behavioural and neuroanatomical findings suggest that young adult mice, particularly 3xTgAD mice, may exhibit resilience to the high-fat diet, limiting the manifestation of deficits. Notably, 3xTgAD mice already show smaller brain volumes at baseline, which may constrain the detectable impact of the diet. Nevertheless, interventions such as exercise appear to support brain structure and function even in the absence of pronounced high-fat diet impairments, potentially by enhancing compensatory or protective mechanisms.

In this work, sex was included as a biological variable in all statistical models. Overall, sex-related effects were most apparent for body weight and behaviour. Males in the combined diet and exercise group showed weight loss, whereas females generally exhibited weight stabilization. Behavioural outcomes also differed by sex, with improvements in spatial memory being more consistently observed in males, while females showed more variable responses depending on genotype and intervention. By contrast, neuroanatomical measures were less sensitive to sex, as neither region-based volumetry nor voxel-wise DBM revealed significant sex differences after correction for multiple comparisons. Together, these findings indicate that, within the age range examined, sex contributes to differences in some of the responses to interventions, whereas structural brain changes appear largely similar between males and females.

Using a multivariate approach (PLS), we identified a brain pattern characterized by positive loadings in the cerebellum and hippocampus, which negatively correlated with a high-fat diet and positively correlated with physical activity and diet interventions. Importantly, this pattern was delineated without relying on predefined neuroanatomical boundaries, demonstrating the power of data-driven methods to capture complex interdependencies across thousands of voxels. Nevertheless, the interpretation of patterns derived from dimensionality reduction approaches remains an active area of discussion (Danyluik et al., 2024; Liu et al., 2022; Tang et al., 2022). Work from our laboratory (Danyluik et al., 2024) suggests that the primary latent variable can often appear significant due to the inherent permissiveness of the Procrustes-rotated permutation test. To ensure the patterns reported here are robust for interpretation rather than a statistical artifact, we integrated the significance along with the latent variables’ strength and their spatial stability through bootstrapping, as evidenced by the high voxel-wise bootstrap ratios in the cerebellum and hippocampus. The convergence of these metrics suggests that the patterns found here encode meaningful neurobiological features that may help us find the underlying biological mechanisms, as exposed by our further SGEA analysis.

Previous work correlating spatial neuroanatomical changes to spatial patterns of gene expression data found that changes in neuroanatomy can help pinpoint genes responsible for the observed phenotypes (Fernandes et al., 2017). Here, the SGEA analysis correlating gene expression data to the PLS-LV3 brain pattern uncovers GO terms associated with biological processes such as glucose homeostasis and positive regulation of fatty acid transport. Here, five out of the seven significant terms are related to glucose homeostasis. These processes have been discovered through other studies, for example, a study in a transgenic model for AD (and WT counterpart) found that a low-fat diet reversed the alterations in glucose metabolism in mice fed a high-fat diet, taking glucose to comparable levels as control mice (fed with a low-fat diet) (Walker et al., 2017). Another study found that a high-fat diet enhanced glucose intolerance in a transgenic model of AD, reversible with the injection of insulin (Vandal et al., 2014). We also found an association with a negative regulation of the response to oxidative stress. In work from Sah et al., a high-fat diet exacerbated oxidative stress in 3xTg-AD mice by inactivating Akt-Nrf2 signaling and reducing downstream antioxidant enzymes, leading to neuronal apoptosis and early cognitive decline (Sah et al., 2017). Lastly, it has been reported that exercise could promote regulation of oxidative stress and regulation of fatty acid transport (Camiletti-Moirón et al., 2013; Liśkiewicz et al., 2020; Souza et al., 2022). For example, a study found that exercise decreased reactive oxygen species in rats fed with a high-fat diet (Molteni et al., 2004).

The data we gathered indicates a potential function of the cerebellum that merits further investigation. Traditionally, the cerebellum has been associated with motor control; however, new evidence supports that it might also be related to cognitive functions (Koziol et al., 2014). Previous research shows a decrease in the neuronal density in both the cerebellar cortex and deep nuclei of the cerebellum in 3xTgAD mice (Esquerda-Canals et al., 2013). This suggests a genetic vulnerability in 3xTgAD mice, which may contribute to the pronounced cerebellar changes observed. Another study found that APP and apoptosis-related molecules (e.g., caspases) increase in the cerebellum of WT mice after only 7 days on a high-fat diet (Xu et al., 2021). A study on children found that physical fitness was associated with larger volumes in some regions of the cerebellum (Jalanko et al., 2024). Another study showed that children who reported difficulty controlling how much they eat had lower grey matter in the parahippocampal gyrus and the left cerebellar lobule VI (Pearce et al., 2023). It has been proposed that since eating behaviours are influenced by predictions (e.g., hunger before a meal, or expectations of food taste) and feedback (e.g., satiety after eating), the cerebellum might be involved in managing prediction errors in these domains, adjusting responses to align with sensory feedback and expectations across motor, cognitive, and affective processes (Iosif et al., 2023).

In terms of limitations, our study used a high-fat diet in which 60% of calories were derived from fat. While high-fat diets for mice typically range between 40–60% of calories from fat, with varying compositions of saturated and unsaturated fatty acids (Lai et al., 2014), this level substantially exceeds the fat intake commonly observed in humans. For example, in the U.S., fat accounts for approximately 30-40% of total calories (Ford and Dietz, 2013), and the most recent report from Canada (2000) indicates a similar intake of around 30% (Gray-Donald et al., 2000). Consequently, the effects observed in our animal model may not fully reflect human dietary patterns. Another limitation is that we used young adult mice (up to 6 months old), corresponding to early adulthood in humans (Cottam et al., 2025). This restricts the generalizability of our findings, as resilience and susceptibility to interventions may change with age. Nevertheless, studying early-life exposures in preclinical models remains valuable, as it can provide insights into mechanisms and potential intervention strategies during the extended preclinical phase of dementia. Additionally, our findings related to the biological processes behind interventions should be confirmed. Future work should address this question as well as the best timing for the initiation of these interventions. Finally, we acknowledge that the use of non-littermate WT controls may introduce confounds such as gut microbiome heterogeneity (McCoy et al., 2017). However, because the primary aim of this study was to evaluate the effects of a high-fat diet and subsequent interventions, we utilized intra-genotype comparisons. This approach reduces the influence of baseline differences between WT and 3xTgAD on the interpretation of treatment effects; however, such baseline variability should be considered when interpreting genotype effects (comparisons between WT_Control and Tg_Control).

In summary, our findings underscore the potential of diet and exercise interventions to counteract the adverse effects of a high-fat diet on the brain, particularly in the cerebellum and hippocampus. Additionally, they highlight the necessity for more research to investigate the underlying mechanisms and the applicability of these findings to human health.

## Materials and Methods

### Animals

All animal experiments were carried out following the Canadian Council on Animal Care and approved by the McGill University Animal Care Committee (Montreal, QC, Canada). Male and female homozygous 3xTgAD mice (https://www.jax.org/strain/004807, RRID: MMRRC_034830-JAX) and non-transgenic Wild-Type (WT) B6129SF2J mice (https://www.jax.org/strain/101045, RRID: IMSR_JAX:101045) were acquired through the Mutant Mouse Resource and Research Center (MMRRC) and were bred on an F2 hybrid cross between C57BL/6J females (B6) and 129S1/SvImJ males (129S). The 3xTgAD mice have a knocked-in PS1M146V mutation at the mouse PSI locus and two human transgenes: APPSwe and tauP301L (Oddo et al., 2003). The PS1M146V and APPSwe mutations affect the processing of the amyloid-beta precursor protein (APP) into Amyloid-beta (Duff et al., 1996), while TauP301L induces tauopathy (Lewis et al., 2000). It has been reported that 3xTgAD exhibits cognitive impairments related to memory retention at four months of age, accumulating intraneuronal amyloid plaques in the cortex, hippocampus, and amygdala (Billings et al., 2005). Other studies have reported cognitive impairment, amyloid plaques, and neurofibrillary tangles in the hippocampus at six months of age (Belfiore et al., 2019). A recent characterization of 3xTgAD mice reported a shift in timing in plaque and tangles appearance, with a substantial impact on females due to estrogen-regulated gene promoters (Javonillo et al., 2022; Sadleir et al., 2015). Mice were bred in an in-house colony, and their offspring were weaned during the fourth week after birth. The offspring were housed in groups of three to four mice per cage at the Douglas Mental Health University Institute’s Animal Facility (Montreal, QC, Canada) in standard housing conditions (12h light/dark cycle, with lights on at 8:00 am). Mice received ad libitum access to food and water at all times.

### Experimental design

All mice were on a standard diet from weaning until two months of age (2016 Teklad Global 16% Protein Rodent Diet, Inotiv, Madison, Wisconsin, USA), containing 4% fat from soybean oil, formulated for growth and maintenance. At two months, 80% of both 3xTgAD and WT mice were randomly assigned to an obesity-inducing high-fat diet (Teklad high-fat diet TD.06414 with 60% of kcal from fat, primarily lard, Inotiv, Madison, Wisconsin, USA), while the remaining mice were transitioned from standard diet to the low-fat control diet (Teklad Purified Control Diet TD.08806 with 10.5% of kcal from fat, primarily soybean oil and lard, Inotiv, Madison, Wisconsin, USA). The control diet is formulated by the manufacturer as the appropriate ingredient-matched control for the high-fat diet, differing only in its macronutrient composition but matched for ingredients and processing. This approach helps isolate the effects of fat content, minimizing confounding variables related to food composition or manufacturer. For four to six months, mice fed a low-fat diet continued on it and served as controls (groups: WT_Control and Tg_Control), while a subset of mice on the high-fat diet remained on it (WT_Highfat and Tg_Highfat). During this same period (4-6 months), to assess the reversibility of the induced obesity, the rest of the mice fed with the high-fat diet were randomly assigned to either a low-fat diet intervention (groups: WT_Diet and Tg_Diet), physical activity while still on the high-fat diet (groups: WT_Exercise and Tg_Exercise) or the combination of exercise and a low-fat diet (groups: WT_Both and Tg_Both). In the physical activity groups, mice had access to a running wheel for voluntary physical activity. Food intake was not monitored due to the difficulties in obtaining accurate measurements as a result of communal housing (Starr and Saito, 2012). Similarly, mice were not placed in isolation to accurately measure wheel use because this introduces the effect of social isolation, which is also a risk factor for AD (Goh and Ladiges, 2015; Huang et al., 2015; Ren et al., 2023). Weight measurements were collected for at least 18 weeks, from two to six months of age (N = 156 mice with 18 measurements each). T1-weighted MRI scans were acquired at two, four, and six months, followed by behavioural testing with the MWM and NOR tests at six months of age. Out of the initial 165 mice involved (around 16 mice per group and a nearly equal number of males and females), 9 died or were sacrificed before the final neuroimaging timepoint, primarily due to fatal injuries as a result of fighting (four males and one female) which was precipitated primarily by the introduction of the running wheel (fighting caused an entire cage of 3xTgAD mice in the combined intervention to be sacrificed), but also from hydrocephalus (one male, one female), and complications as a result of MnCl_2_ injections (which were needed for improve MRI contrast) (two males). Fig 1A shows the experimental workflow, Supplementary Table S3 describes the nutrient information per diet, and Supplementary Table S4 shows the timeline at which each group received their respective diets and the N per group at each scanning time point.

### Magnetic Resonance Image acquisition

T1w MRI images were acquired at three time points, when mice were two, four, and six months old. These time points were selected to capture the neuroanatomy at baseline, before, and after interventions. Scans were collected using a 7T small animal scanner (Bruker Biospec 70/30 USR; 30-cm inner bore diameter) at the Douglas Mental Health University Institute (Montreal, QC, CA). Mice were injected 24 hours before the scanning with a bicine-buffered solution of manganese (II) chloride (MnCl_2_) titrated with 1M sodium hydroxide (NaOH; Sigma Aldrich, Switzerland) at a dosage of 62.5 mg/kg. MnCl_2_ enhances the contrast of T1w structural MRI scans due to the paramagnetism of Mn_2+_ that reduces the T1 relaxation time, enhancing the contrast between tissues ^(^Lee et al., 2005; Yang and Li, 2020^)^. NaOH and bicine were added to mitigate the acidic environment caused by MnCl_2_, preventing tissue damage at the injection site and keeping the pH at about 7.4 (Seo et al., 2011). Dosage was calculated based on weight at baseline to avoid toxic doses in obese mice. High doses were observed to cause skin lesions in previous work from our group (Rollins et al., 2019). The injection was administered to the intraperitoneal cavity to limit toxicity (Kuo et al., 2005). Before the scanning, animals were anesthetized with isoflurane (Fresenius Kabi Canada Ltd, Richmond Hill, CA) at 5% and injected with 0.5 mL of 0.9% sterile sodium chloride physiological solution (Sigma Aldrich). During the scanning, mice were maintained under 1-2% isoflurane, with breathing rate monitored and maintained at 30-100 breaths per minute. The sequence used to acquire the images was a 3-dimensional fast low-angle shot (3D FLASH), and the final T1w MRI was generated from an average of two acquisitions (TR = 20 ms, TE = 4.5 ms, matrix size = 180 x 160 x 90, voxel dimensions = 100 μm isotropic, and flip angle = 20°) and fat suppression. Images were collected with the Bruker 23 mm volumetric transmit/receive quadrature coil (Anastassiadis, 2020).

### Image processing

Raw Digital Imaging and Communications in Medicine (DICOM) scans were converted to the Medical Imaging Net CDF (MINC) file format and preprocessed with a set of tools and libraries developed in our laboratory and at the Montreal Neurological Institute (MNI), freely available online (Minc-toolkit 1.9.16, RRID: SCR_014138). Preprocessing included stripping images of their native coordinate system, left-right flipping of images to compensate for the scanner’s native radiological coordinate system, denoising using a patch-based non-local means filter previously validated (Coupe et al., 2008), and affine registration to an average mouse template to produce a brain mask. N4ITK was used as an improved version of the N3 bias field correction algorithm for inhomogeneity correction at a minimum distance of 5 mm (Tustison et al., 2010). Then, a de-scaled and de-sheared version of the affine registration was used to perform a rigid alignment of the scan to the mouse template space (Gallino et al., 2019). Images were inspected using the Display visualization software of MINC tools. Quality control was performed for each time point.

The inspection was mostly focused on detecting the presence of hydrocephalus or motion-related artifacts. Of the 165 scans collected at timepoint one, 160 passed quality control (https://github.com/CoBrALab/documentation/wiki/Mouse-QC-Manual-(Structural)); from the 160 scans collected at timepoint two, 148 passed; and from the 156 scans collected at timepoint three, 145 passed quality control. Therefore, we used 453 brain images for further analysis.

### Volumetric analysis

We performed a volumetric analysis on mice with at least one image that passed quality control (total N = 161 mice, 453 scans) to study whether the high-fat diet and intervention strategies impacted the neuroanatomy of 3xTgAD and wild-type mice. We parcellated the brain into 182 structures per hemisphere according to the DSURQE (Dorr-Steadman-Ullmann-Richards-Qiu-Egan) mouse brain atlas (Dorr et al., 2008; Richards et al., 2011; Steadman et al., 2014; Ullmann et al., 2013) using the Multiple Automatically Generated Templates (MAGeT) brain segmentation algorithm (Chakravarty et al., 2013). MAGeT first performs a non-linear transformation between the input data (T1w MRI scans) and the input atlas. The atlas is registered to a small subset of the subject images (n=21). These labeled images are used as a template library, and pairwise non-linear transformation is performed to generate as many segmentations per subject as there are templates in the library (n=21). Given there are now 21 ‘atlases’, these brains are now registered to each subject such that the final segmentation is produced by a voting procedure at the voxel level, where the most voted label is selected. Quality control of the outputs was performed by visually inspecting each segmentation overlaid on each mouse scan and verifying that the labels matched the brain anatomy (https://github.com/CoBrALab/documentation/wiki/MAGeT-Brain-Quality-Control-(QC)-Guide). Out of 453 MAGeT segmentations, 385 passed quality control, corresponding to scans from 157 mice: 139 scans from the first timepoint, 116 scans from the second, and 130 scans from the third. Due to frequent artifacts in the olfactory bulbs, labels from this region were excluded from the volumetric analysis (see Supplementary Table S5). Importantly, these artifacts did not impact the voxel-wise analysis detailed in the following section. Volumes were computed from these segmentations based on the number of voxels labelled for each region and the voxel dimensions.

### Deformation-based morphometry analysis

After the volumetric study, we aimed to analyze the changes in volume without any *a priori* assumptions on neuroanatomical boundaries. We applied a deformation-based analysis pipeline to model voxel-wise volume changes using the Jacobian determinant, which measures voxel contraction or expansion relative to a template (Chung et al., 2001; Lau et al., 2008). Images from mice with at least two time points that passed quality control (N = 152 mice, 423 images) served as input. To study overall brain differences, absolute Jacobians modeling the linear and non-linear transformations were generated using the two-level model-building toolkit Pydpiper (Friedel et al., 2014). Images were non-linearly registered to create a population-specific average template, followed by a group-wise registration to account for the longitudinal nature of our data (Friedel et al., 2014). The non-linear registration was performed using the diffeomorphic MINC Advanced Normalization Tools (mincANTs) algorithm (Avants et al., 2011, 2009). The absolute Jacobians were log-transformed and blurred (0.2 mm full-width at half-maximum) to conform to Gaussian assumptions for statistical testing (Chung et al., 2001; Gallino et al., 2019). For within-subject registration, an average representation of each specific subject’s anatomy over time was created using a hierarchical multi-resolution registration strategy (Avants et al., 2009). These subject-specific averages served as inputs to build the group average and the common space for group-wise and longitudinal comparisons (Gallino et al., 2019; Kong et al., 2018). Resampled absolute Jacobians adjusted for whole-brain size were used for all analyses. Jacobian determinants quantify local volumetric differences relative to the population average, such that values greater than 1 indicate local expansion (relative to the population reference) and values less than 1 indicate local contraction; values equal to 1 indicate no change. For voxel-wise longitudinal examination of volume over time between groups, a group-specific baseline mean was calculated for each group, such that changes could be interpreted as deviations from baseline over time. These baseline-centered values were subsequently exponentiated and expressed as volume percentage change from baseline for improved visualization and interpretation.

### Partial Least Squares analysis

To study the relationship between voxel-wise neuroanatomical changes and behavioural data, we ran a behaviour Partial Least Squares analysis (PLS) in Matlab 2022a (PLScmd software; Baycrest, Canada) as previously done in other work from our group (Guma et al., 2021). PLS can be used when the number of features (e.g., voxel-wise data) is higher than the number of observations (e.g., number of mice), despite the presence of correlated variables (collinearity) (Cramer, 1993). We constructed a matrix (X) made of voxel-wise Jacobian values from the last MRI scan collected (third timepoint, N = 96 mice). Another matrix (Y) was built with data from 45 variables, including demographics and behavioural data from the MWM and NOR tests. The matrices X and Y were correlated and subjected to a mathematical process (singular value decomposition) to obtain latent variables (LV). Only mice with complete MRI and behavioural data were included; this led to the exclusion of mice in the groups that received the combined intervention (Tg_Both and WT_Both). PLS produces as many LVs as there are variables in the behavioural data, in our case, 45. LVs capture the maximal shared variation between the two data sets, summarizing their relationship in a single dimension, and can be interpreted as maximally covarying brain patterns and their associated behavioural factors (McIntosh et al., 1996). Behavioural variables used for partial least squares analysis were: 1) Demographic: sex, genotype, low-fat or high-fat diet, intervention type, no intervention; 2) Variables from Morris Water Maze: total path in meters, total time, time in quadrant of interest, time and distance in other quadrants, speed, count of passthroughs above the platform, passthroughs near the platform zone, GCM, cumulative time in the platform zone; 3) Variables from the novel object recognition test: time in the boundary of the novel object (seconds), time in the boundary of the familiar object (seconds), percentage of time of exploration of the novel object, preference for novelty, frequency in the novel object exploration, percentage of time of exploration in the boundary of the novel object, percentage of time of exploration in the boundary of the familiar object, distance (meters), velocity, frequency entering the arena, time spent in the arena (seconds), frequency entering the zone of the novel object, frequency entering the zone of the familiar object, time spent in the zone of the novel object (seconds), time spent in the zone of the familiar object (seconds), frequency entering the zone of the boundary of the novel object, frequency entering the zone of the boundary of the familiar object, time spent in the zone of the boundary of the novel object, time spent in the zone of the boundary of the familiar object, frequency of high-mobility, cumulative time of high-mobility, frequency in mobility, cumulative time of mobility, frequency in immobility, cumulative time immobile, moving time, not moving time.

### Spatial Gene Enrichment analysis

To assess the biological mechanisms underlying the brain pattern identified using PLS (latent variable 3), we conducted a Spatial Gene Enrichment Analysis (SGEA) in R/4.1.3. We utilized gene expression data from the Allen Institute’s Mouse Brain Atlas (Lein et al., 2007), which contains 4345 three-dimensional images of gene expression for 4082 unique genes acquired by immunohistochemistry on brain slices from adult (P56) male C57Bl/6 mice. Additionally, we used the high-resolution Allen Mouse Brain Common Coordinate Framework template (CCFv3, 200 µm voxel size) (Wang et al., 2020) to provide a standardized anatomical reference space, allowing gene expression maps to be aligned and compared consistently across the brain. Gene annotations, which link individual genes to specific biological processes (also called modules), were drawn from the Gene Ontology (GO) consortium. We used the gene annotation dataset MOUSE_GO_bp_no_GO_iea_symbol, with 15757 gene sets (Gene Ontology Consortium, 2015). These annotations allow us to group genes into functional modules, providing biological context for interpreting patterns of spatial gene expression. The annotations used here were inferred from electronic evidence, meaning they were computationally predicted rather than manually curated. The gene expression images and CCFv3 template were realigned to RAS+ orientation and converted to the MINC format. Next, we conducted the SGEA analysis in R/4.1.3 using the bootstrap ratio map from PLS latent variable 3 (LV3) as input. The map was thresholded at the bootstrap ratio value of 1.95 (95% confidence interval) to include only regions contributing significantly to LV3, and are therefore meaningfully related to behavioural measures. To explore the molecular mechanisms underlying these patterns, the thresholded brain loading map was spatially correlated with gene expression profiles. Genes were then ranked based on the strength of their correlation with the brain pattern using Spearman correlation.

### Behavioural tests

Novel Object Recognition (NOR) was performed one week after the third MRI scan, and the Morris water maze (MWM) test was performed two days after the NOR test. For the NOR test, on day zero, mice were placed in a cubic open field box (45 cm^3^) with bedding from their cage. Mice were allowed to explore for 10 min. On day one, mice were placed in the same box with bedding from their home cage, and they were allowed to explore two identical objects (4 cm^3^ each) for 10 min. The objects were spaced 20 cm apart. Mice were returned to their home cage, and one hour later, they returned to the test box with one object replaced with the novel object. Mice were given 10 minutes to explore. The location of the novel object was randomized (left or right side) and balanced across trials. The tests were conducted under a red light with an overhead camera. The camera recorded the test, and the resultant video clips were converted from DAV to AVI and further edited to last precisely 10 minutes. Videos were analyzed with Ethovision XT 12 Software. Only data from the test trial was used. During analysis, a boundary area around the object was set to 10 cm². To determine if mice spent more time around the novel object, the percentage of exploration in the boundary zone of the novel object was calculated. To do so, the time spent in the boundary area of the novel object was divided by the sum of the time spent in the boundary area of the novel and the familiar object and multiplied by 100. Another variable of interest was the discrimination index, also known as the discrimination ratio (Antunes and Biala, 2012). If positive, the discrimination ratio indicates more time spent exploring the novel object. A value of zero means equal time spent on both objects. This variable was calculated as the amount of time spent exploring the boundary zone of the novel object minus the amount of time spent exploring the boundary zone of the familiar object, divided by the total amount of time spent exploring the boundary zone of both objects.

The MWM test was performed in a circular tank (120 cm in diameter) with water at 21-22 ℃. Water was made opaque by adding white, non-toxic tempera paint. There were three visual cues on the interior wall of the tank above the water line (in the north, east, and southwest). An escape platform with a diameter of 15 cm was used. On day zero, mice were habituated with a trial where the escape platform was visible (1 cm above the water line). The anterior allowed the identification of apparent sensorimotor deficits. From day one to five, mice were trained five times by entering the tank with a non-visible escape platform always located in the southwest quadrant. During this training, mice entered the tank from one of three starting points (northeast, southwest, and southeast) in pseudorandom order. If the mice couldn’t find the platform in 60 seconds, they were gently guided towards it. A failure was a training in which mice didn’t find the platform within 60 seconds. On day six (probe day), a single trial was performed. In this trial, the escape platform was removed. Mice entered through the northeast point and were allowed 60 seconds to explore. Each trial was recorded with a mounted camera above the tank. However, some mice were recorded with a different camera due to software failure. Only data from the probe day was used. Mice were excluded if they failed to find the platform in three of six training sessions. The data from the first camera was analyzed with tracking software (HVS Image Ltd). The data captured with the second camera was analyzed using a different software (Ethovision Noldus XT 12). Variables of interest were further analyzed, such as the average cumulative distance from the platform (also known as cumulative Gallagher’s measure) and the number of pass-throughs (passages above the previous platform location). The cumulative distance from the platform is recognized as a solid measure of memory, and it has a good sensitivity to small differences in groups (Pereira and Burwell, 2015; Vorhees and Williams, 2006).

## Statistical Analyses

### Weight and behaviour

Body weight data were analyzed with Linear Mixed Effects (LME) models implemented in the lme4 package in R (version 4.1.3). The model included a four-way interaction between time (in weeks), intervention, genotype, and sex as fixed effects, with a random intercept and slope for weeks at the individual level to account for repeated measurements within each mouse (*Weight* ∼ *Week*\**Intervention*\**Genotype*\**Sex* + (*Week*|*ID*)).

Because higher-order interaction terms are difficult to interpret directly (Busenbark et al., 2022), we estimated model-based rates of weight change over time using the ‘marginaleffects’ package (version 0.28.0). These estimates (referred to as marginal comparisons) quantify the expected change in body weight per week (g/week) for each combination of intervention, genotype, and sex. Ninety-five percent confidence intervals (95% CI) were computed using the delta method as implemented in ‘marginaleffects’ (Arel-Bundock, 2025).

Behavioural data derived from the MWM and NOR were statistically assessed using regression models selected based on the properties of each variable. For each outcome, data distributions were visually inspected using histograms (available in Supplementary Material, Fig S7) and formally evaluated for normality within experimental groups using the Shapiro-Wilk test and for homogeneity of variance using Levene’s test. No outliers were detected for MWM; however, three outliers were removed due to extreme values for NOR (One mouse from the WT_HighFat group and two mice from the Tg_HighFat group). Model families were chosen according to the outcome, for example, beta regression for bounded proportion outcomes, generalized linear models for count data, and hurdle models for zero-inflated outcomes. A detailed summary of these choices is available in Supplementary Material (Tables S1-2). For NOR, two competing models (*Behavioural Outcome* ∼ *Intervention* * *Genotype* * *Sex* or *Behavioural Outcome* ∼ *Intervention* * *Genotype* + *Sex*) were compared using model performance metrics within the ‘model_performance’ function from the ‘performance’ package (version 0.15.2) in R (version 4.4.3) to decide which model best fits the behavioural outcome being analyzed. For MWM, the same two competing models were compared, with additional covariates for *Software* (see Materials and Methods, section ‘Behavioural tests’ for more details) and *Weight* (*Behavioural Outcome* ∼ *Intervention* * *Genotype* * *Sex + Software + Weight* or *Behavioural Outcome* ∼ *Intervention* * *Genotype* + *Sex + Software + Weight*). To interpret significant interaction terms, model-based marginal contrasts were computed using the ‘contrast’ function from the ‘marginaleffects’ package. Marginal contrasts quantify the differences between each intervention and the Control (mice that remained on a low-fat diet throughout the experiment), while holding all other covariates at fixed values. These effects can be interpreted as the expected change in the outcome attributable to the intervention, independent of other variables in the model. Only mice with full behavioural data were included in the analyses. The number of mice per group for NOR and MWM is provided in Supplementary Table S6. Results were plotted using ggplot2 in R.

### Volumetric and deformation-based analyses

The association between volume data derived from MAGeT and groups was evaluated with a hierarchical LME model with the RMINC/v1.5.3.0 package for R (Lerch et al., 2017). The model included volume per region as the dependent variable, an interaction between group and age in days, sex as an independent variable, and a random intercept allowed per mouse (*Volume* ∼ *Group*\*poly(*Age*,2) + *Sex* + (*1|ID*)). The poly(*Age*,2) term allows the model to capture quadratic (second-degree polynomial) relationships between Age in days and volume while considering interaction effects with each group; data from the three time points were included. The linear mixed effects model output was corrected for multiple comparisons with False Discovery Rate (FDR) with a 10% threshold (Benjamini and Yekutieli, 2001), meaning that among the regions shown as significant, we expect at most 10% to be false positives. Results were plotted by overlaying the statistics map on top of the population average using functions available in RMINC. The plots display the linear terms of the interaction between group and age in days (*Group*\*poly(*Age*, 2)[1]), since the quadratic interaction didn’t yield any significant results after FDR correction.

In the case of the deformation-based morphometry analysis, we assessed statistical significance by fitting an LME model at each voxel using the RMINC/v1.5.3.0 package for R (Lerch et al., 2017). The model used baseline-centered log-transformed absolute Jacobian determinants as the dependent variable, an interaction term between group and age in days (modeled as a quadratic polynomial), sex as an independent variable, and a random intercept allowed per mouse (*baseline-centered log-Jacobian* ∼ *Group*\*poly(*Age*,2) + *Sex* + (*1|ID*)). Centering was performed by subtracting the group-specific baseline mean, such that model estimates represent deviations from baseline. Data from the three time points were included. The model outputs were corrected for multiple comparisons with False Discovery Rate (FDR), with results being displayed with a 10% threshold, meaning that among the voxels shown as significant, <10% will be false positives. Results were plotted by overlaying the statistics map on top of the population average. Visualization of voxel-wise trajectories of peak voxels was done using RMINC and ggplot2. As in the case of the volumetric results, the plots display the linear terms of the interaction between group and age in days (*Group*\*poly(*Age*, 2)[1]), as the quadratic interaction didn’t yield any significant results.

### Partial Least Squares Analysis

The output LVs were statistically assessed by permutation testing (3000 tests), and the contribution of individual variables to each LV was assessed with bootstrap resampling to estimate standard errors and confidence intervals (3000 samples) (McIntosh and Lobaugh, 2004). Bootstrap is a statistical resampling technique that creates several samples (with replacement) from the original dataset to assess the variability of the analysis. The bootstrap ratio (BSR) is calculated as the original weight/loading derived from the PLS, divided by the standard error of the weights or loadings obtained from the bootstrap resampling. A BSR is frequently compared with a standard normal distribution (e.g., a threshold of 1.95 for a 95% confidence interval) to assess significance. BSRs are used to determine which brain areas significantly relate to behavioural measures within an LV.

### Spatial Gene Enrichment analysis

For the SGEA analysis, we used the tmodUtest function (tmod/0.46.2 package in R) to assess module enrichment. Genes were ranked based on their spatial correlation with the brain phenotype (PLS-LV3 loading map) using Spearman correlation. Gene modules (groups of genes with shared biological functions) were filtered to include only genes that were present in the gene expression data from the Allen Institute’s Mouse Brain Atlas that we had available. Modules were further restricted to contain between 10 and 500 genes. For each module, we determined which of its genes appeared in the ranked list and where they fell in the ranking. This step allowed us to assess whether certain modules are enriched with highly ranked genes (i.e., genes that are strongly correlated with the phenotype). The Mann-Whitney test was used to compute a U-statistic and p-values for the ranked gene list, with further transformation to the Area Under the Curve (AUC) to measure effect size. The significance of the AUC was tested by comparing against randomized AUC null distributions (Fulcher et al., 2021), with p-values adjusted using FDR correction.

## Funding

Healthy Brains Healthy Lives (CLG)

Fonds de recherche du Québec - Santé (CLG, CA, MU)

## Author contributions

Conceptualization: CA, MMC

Data curation: CA

Formal analysis: CLG, CA

Investigation: CA, MP, MU, DG

Resources: MMC

Software: GAD, YY

Supervision: MMC Visualization: CLG

Writing - original draft: CLG

Writing - review & editing: CLG, CA, MU, DG, ST, GAD, MMC

## Data and materials availability

The data that support the findings of this study are openly available in Zenodo at https://doi.org/10.5281/zenodo.14907093. The repository includes raw data, scripts for the main analyses, and statistical maps generated in this study.

