## Supplementary for "Examining Alzheimer’s Disease modifiable risk factors: Impact of physical activity and diet on neuroanatomy and behaviour in mouse models"

### Supplementary Material

**Content:**


- Supplementary Results, ‘Volumetric Analysis’ (with Fig. S1, S2, and S3)
- Fig. S4. Age-dependent group differences in brain volume
- Fig. S5. Trajectories of significant peak voxels identified for the diet intervention
- Fig. S6. Trajectories of significant peak voxels identified for the exercise intervention
- Fig. S7. Distribution of Behavioral Outcomes
- Table S1. Model selection per outcome in NOR
- Table S2. Model selection per outcome in MWM
- Fig. S8. Significance and percentage of covariance explained by each latent variable
- Fig. S9. Behavioral weights and brain loadings for LV1, LV2, and LV4
- Fig. S10. Relationship between Brain scores for LV3 and weight
- Fig. S11. Relationship between Brain scores for LV3 and GCM, and Passthrough
- Fig. S12. Brain scores and medians per group for LV1, 2 and 4
- Table S3. Nutrient information per diet
- Table S4. Number of mice per time point
- Table S5. List of regions (labels) from the olfactory bulb removed from the volumetric analysis
- Table S6. Number of mice per group for NOR and MWM

### Supplementary Results

#### Volumetric Analysis

We used a hierarchical linear mixed-effects (LME) model to analyze the effects of genotype, high-fat diet, and interventions on brain anatomy across 182 regional brain volumes derived from the MAGeT algorithm with the DSURQE mouse brain atlas. The analysis included scans from 2, 4, and 6 months of age, modeling age as a quadratic term to capture non-linear volumetric changes. The model accounted for group-age interactions, sex, and random intercepts for individual mice. Results were corrected for multiple comparisons using a 10% FDR threshold.

Results in Fig S1A show the linear term for the interaction between group and age in WT mice only, since no significant effects were found on 3xTgAD mice. The quadratic term of the interaction between group and age was not significant for any comparison.

To characterize genotype-related differences in brain volume, we first performed a targeted comparison between 3xTgAD and WT mice within the low-fat diet condition (Tg_Control vs WT_Control), where no dietary manipulation is present. This comparison identified a small but significant cerebellar region (Fig. S2). Given that volumetric differences between 3xTgAD and WT mice are known to be widespread and not restricted to specific interventions, we next examined the main effect of genotype, which captures genotype-related differences that are consistent across age and experimental conditions (Fig. S1A, "Genotype effect"). This analysis revealed significant differences across multiple brain regions and accounts for the global volumetric difference observed in Fig. S1B-C, with 3xTgAD mice showing smaller hippocampal and cerebellar volumes relative to WT mice.

We subsequently assessed the effect of the high-fat diet (Fig. S1A, 'High-fat diet effect'). We found significant effects on cerebellar volumes in WT mice but not in 3xTgAD mice. Finally, we examined the effects of intervention strategies in both genotypes. Again, effects at the volumetric level were found in WT mice only. Physical activity (Fig. S1A, 'Exercise intervention effect') affected regions in the cerebellum, hippocampus, and cortex (<10% FDR), while diet or the combined intervention showed significant effects limited to the cerebellar region (Fig. S1A, 'Diet intervention effect', and 'Effect of combined interventions'). Trajectories for volume changes for the cerebellum and hippocampus (Fig. S1B-C) show that a high-fat diet is related to a volume decline in these structures, compared to groups that received an intervention. This suggests a possible rescue effect caused by intervention strategies in WT mice.

Regarding sex effects, we found significant differences that survived 10% FDR when comparing males versus females (Fig. S3). These differences were found for the contrast between Tg_HighFat and Tg_Control, in the cerebellum, cortex, and caudoputamen. Differences were also present for each intervention in 3xTgAD mice (Tg_Diet, Tg_Exercise, or Tg_Both versus Tg_HighFat), mainly in the cerebellar hemispheres. There were no observed sex differences in WT mice, indicating that the sexes respond differently to all treatments in the 3xTgAD mice.

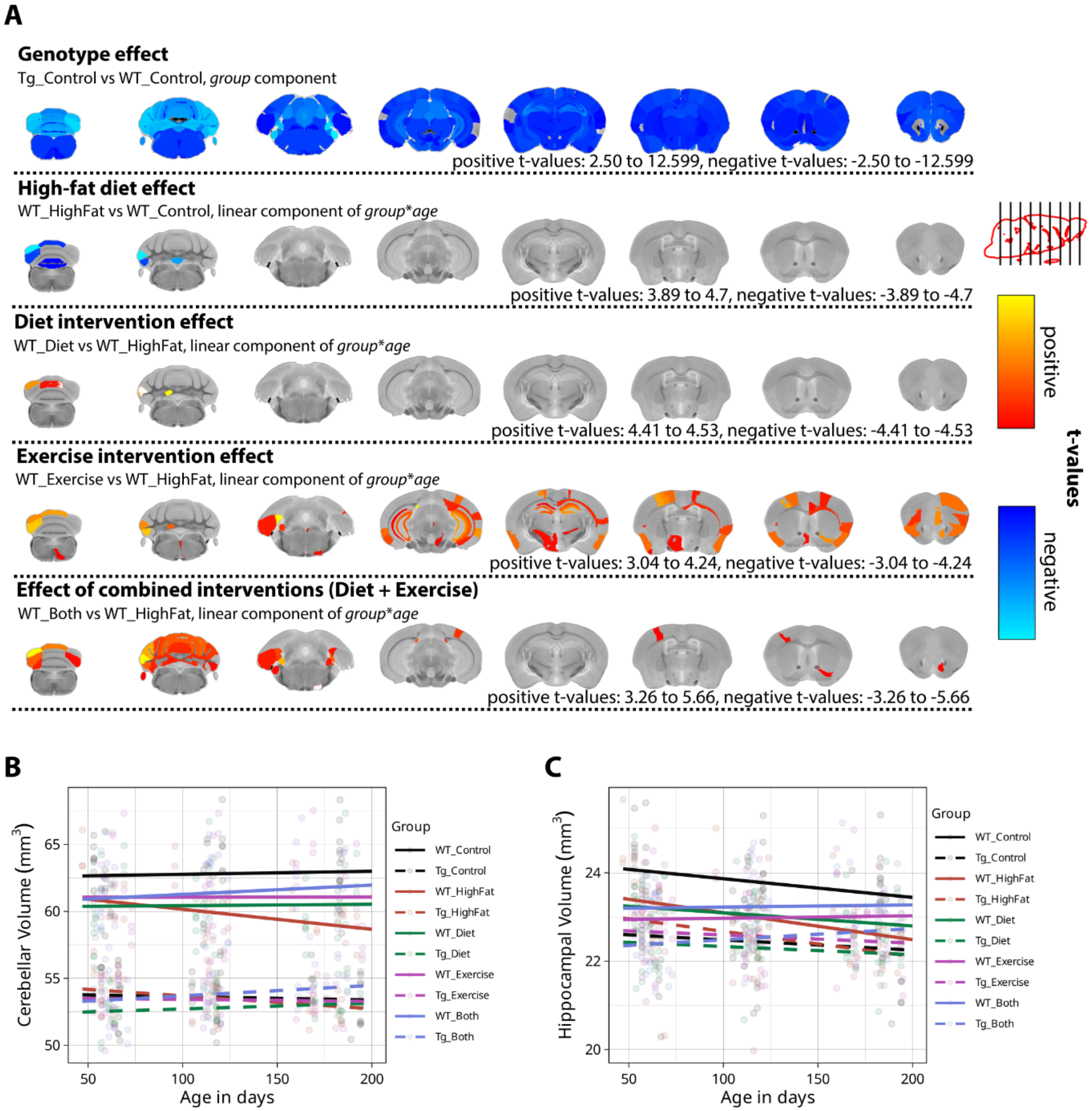


**Fig. S1. Significant volumetric differences.** (A) Statistical maps showing significant differences in brain volume trajectories after false discovery rate correction (FDR, 10%) on top of the DSURQE anatomical atlas. The t-statistic maps show the linear term of the significant interaction between group and age, with the exception of the Genotype effect, which shows the main effect (independent of age). The comparisons are organized into sections based on the main experimental factors: the genotype effect, the high-fat diet effect, and the effect of each intervention strategy. Linear mixed effects model: *Volume* ~ *Group**poly(*Age*,2) + *Sex* + (*1*|*ID*). Regions in blue represent negative t-values, and regions in red represent positive t-values. No significant effects were found on 3xTgAD mice. (B) Cerebellar volume and (C) hippocampal volume in $mm^3$ by group and time point. Solid lines show WT groups and dashed lines show 3xTgAD groups. N = 157 mice. FDR: False discovery rate.

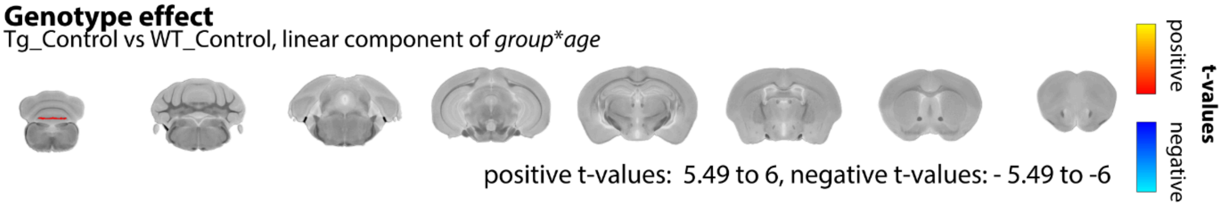


**Fig. S2. Brain volume changes of the group by age in days interaction as determined by volumetric analysis.** T-statistic maps of the linear term of group by age interaction thresholded at 10% FDR overlaid on the anatomical atlas, for the comparison between Tg_Control vs WT_Control groups. Model: *Volume* ~ *Group**poly(*Age*,2) + *Sex* + (*1*|*ID*). N = 152 mice.

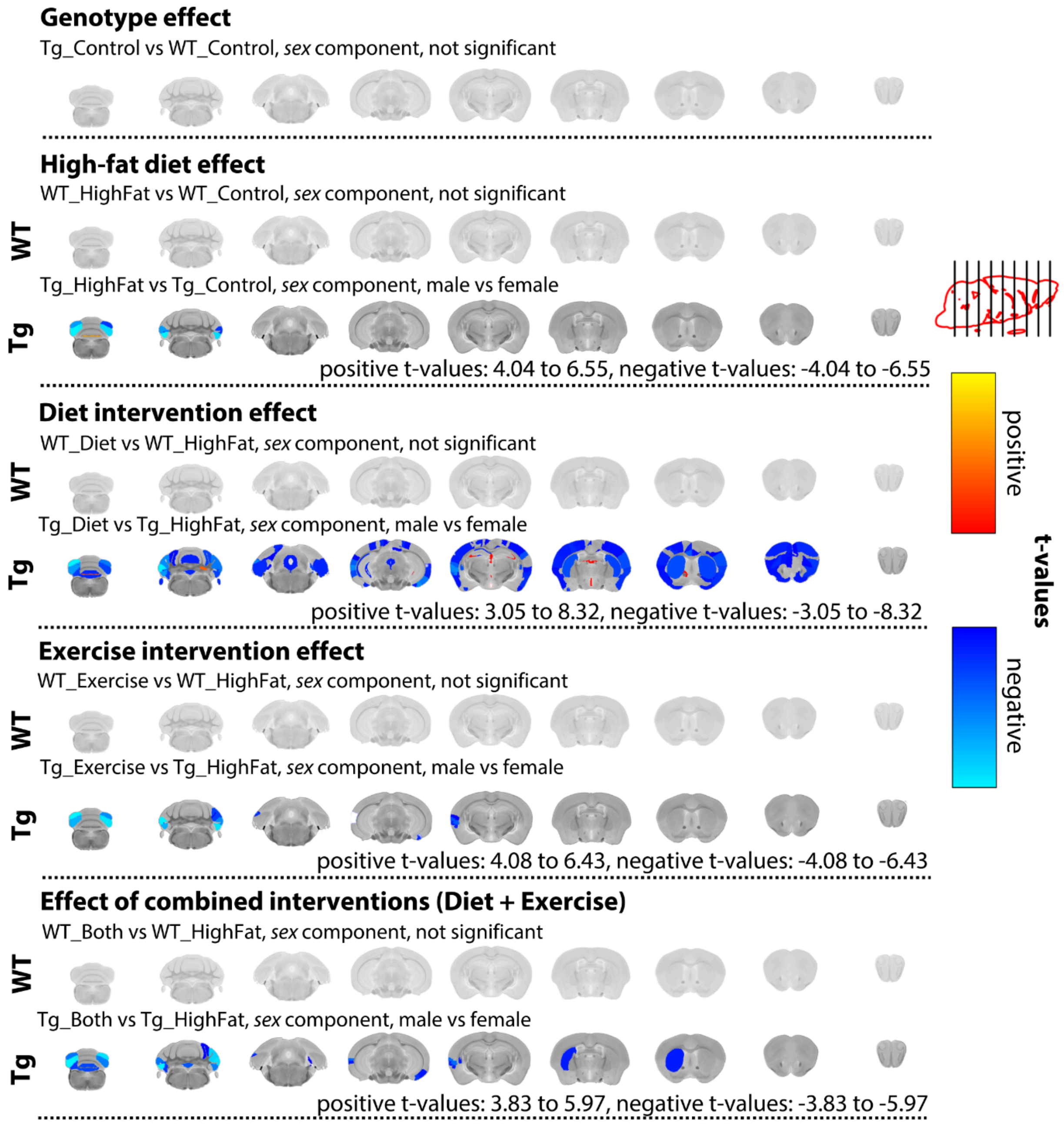


**Fig. S3. Significant sex differences in brain volume trajectories.** Statistical maps showing significant sex differences per contrast after false discovery rate correction (FDR, 10%). T-statistic maps of the significant sex differences are overlaid on top of the anatomical atlas. The comparisons are organized into sections based on the main experimental factors: The genotype effect, the high-fat diet effect on each genotype, and the effect of each intervention strategy on each genotype. N = 157 mice. Regions in blue represent negative t-values, and regions in red represent positive t-values. Model = *Volume* ~ *Group**poly(*Age*,2) + *Sex* + (*1*|*ID*).

***************end of supplementary results****************

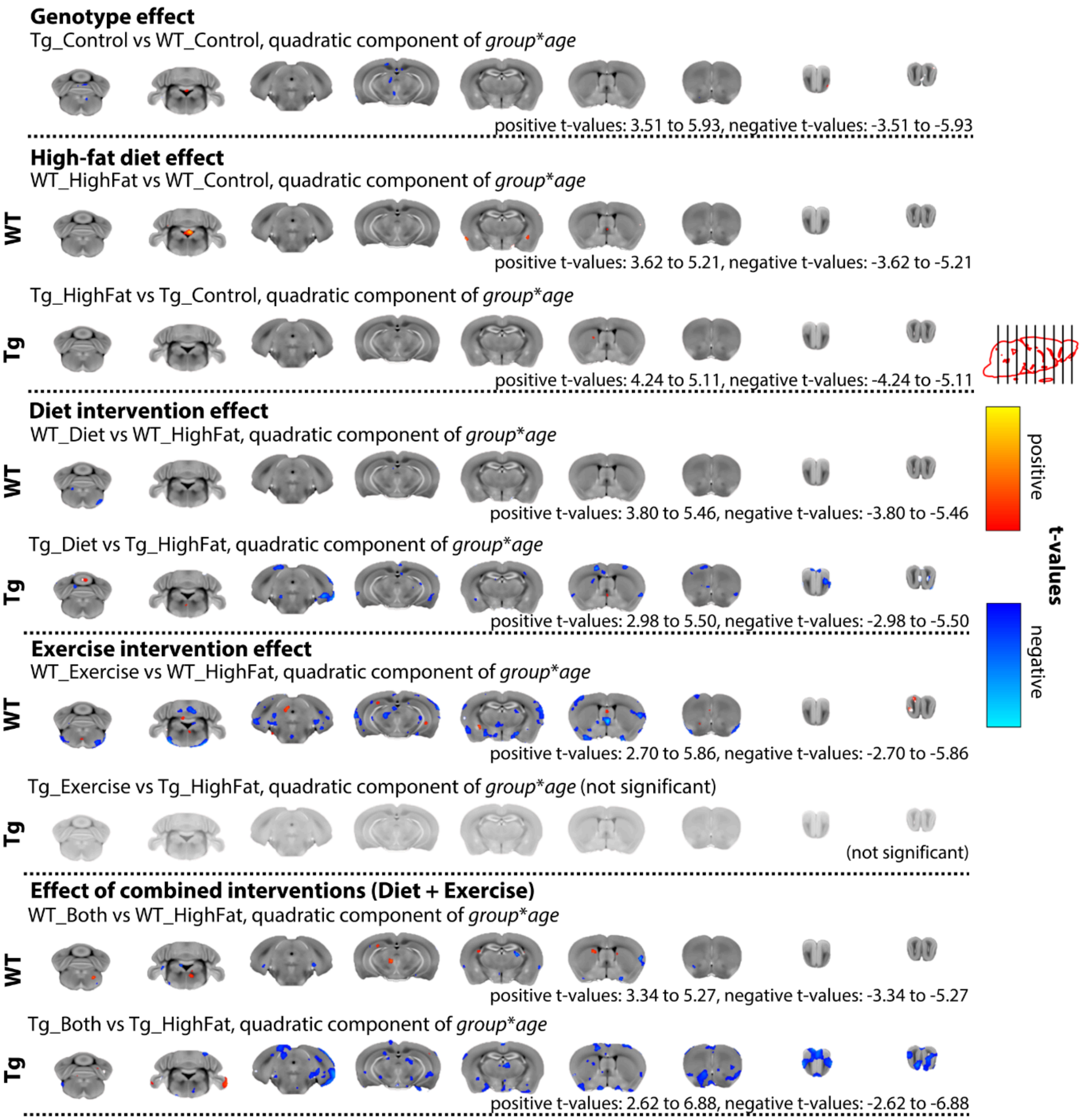


**Fig. S4. Age-dependent group differences in brain volume.** *t*-statistic maps show the quadratic component of the group * age interaction, thresholded at 10% false discovery rate (FDR) and displayed at their peak *t* values, overlaid on the population-average brain. Estimates were derived from a linear mixed-effects model of the log-transformed Jacobian determinant (*Group* × poly(*Age*, 2) + *Sex* + (1 | *ID*)). N = 152 mice.

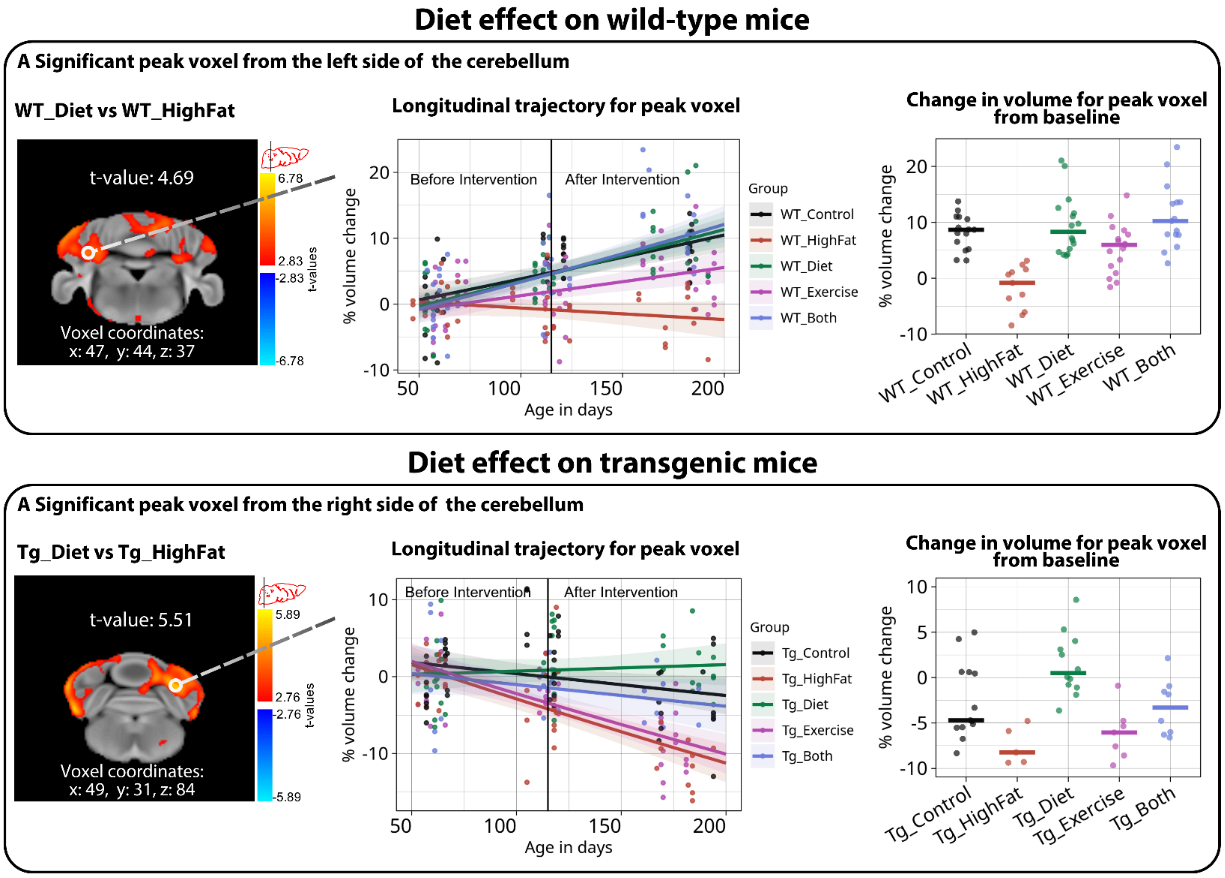


**Fig. S5. Trajectories of significant peak voxels identified for the diet intervention.** Significant peak voxel from the cerebellar region for the comparison between WT_Diet and WT_HighFat (top panel, A). The bottom panel shows a peak voxel selected from the cerebellum from the comparison between Tg_Diet vs. Tg_HighFat (bottom panel, A). The left panels show the statistical maps of the interaction between the group and age in days, thresholded at 10% FDR, t-values, and coordinates from which peak voxels were selected. The middle panels show the percent change in volume from baseline in the selected peak voxels, illustrating volumetric changes across all three time points. The panels on the right show the percent volume change from baseline at the third time point only, highlighting final group differences. FDR: False Discovery Rate.

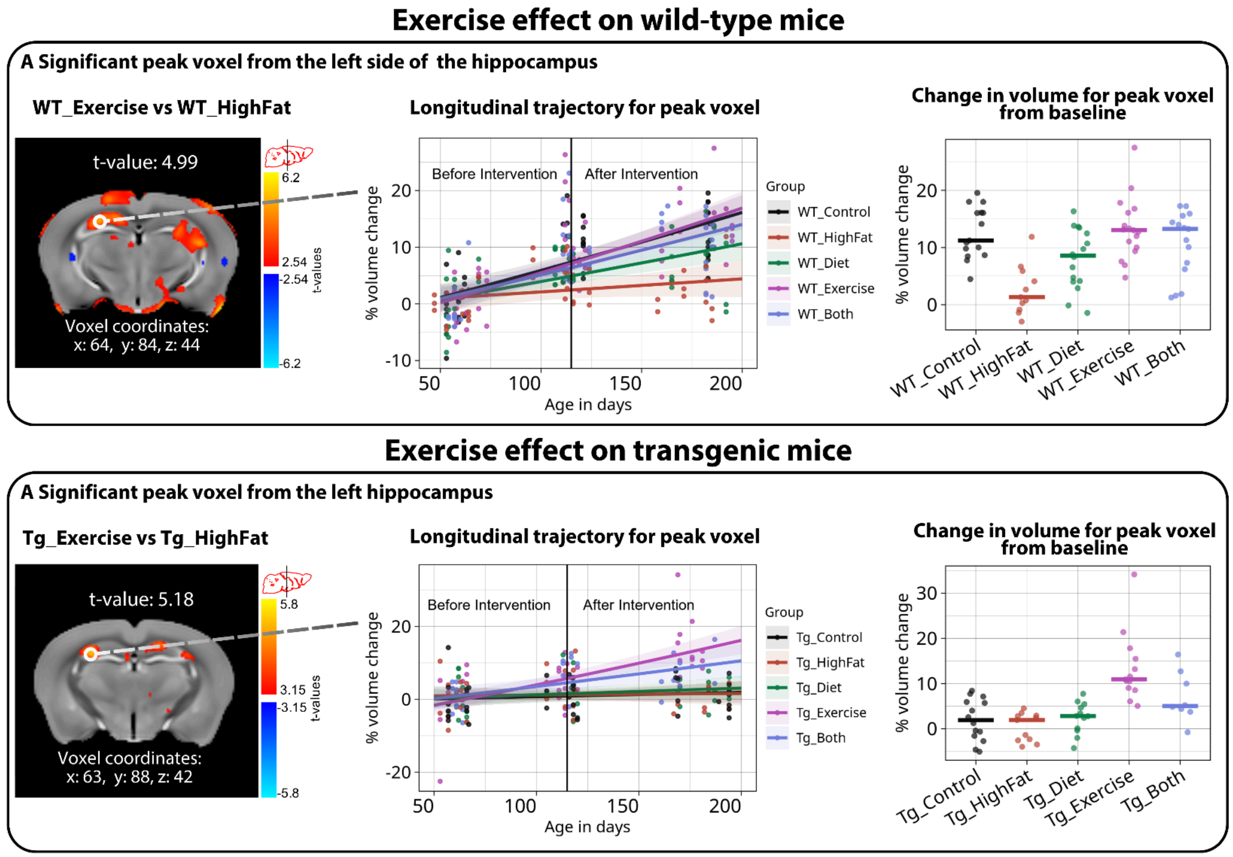


**Fig. S6. Trajectories of significant peak voxels identified for the exercise intervention.** Significant peak voxel from the hippocampal region from the comparison between WT_Exercise and WT_HighFat (top panel, A). The bottom panel shows a peak voxel selected from the left hippocampus from the comparison between Tg_Exercise vs. Tg_HighFat (bottom panel, A). The left panels show the statistical maps of the interaction between the group and age thresholded at 10% FDR, t-values, and coordinates from which peak voxels were selected. The middle panels show percent volume change from baseline in the selected peak voxels, illustrating volumetric changes across all three time points. The panels to the right show percent volume change from baseline at the third time point only, highlighting final group differences. FDR: False Discovery Rate.

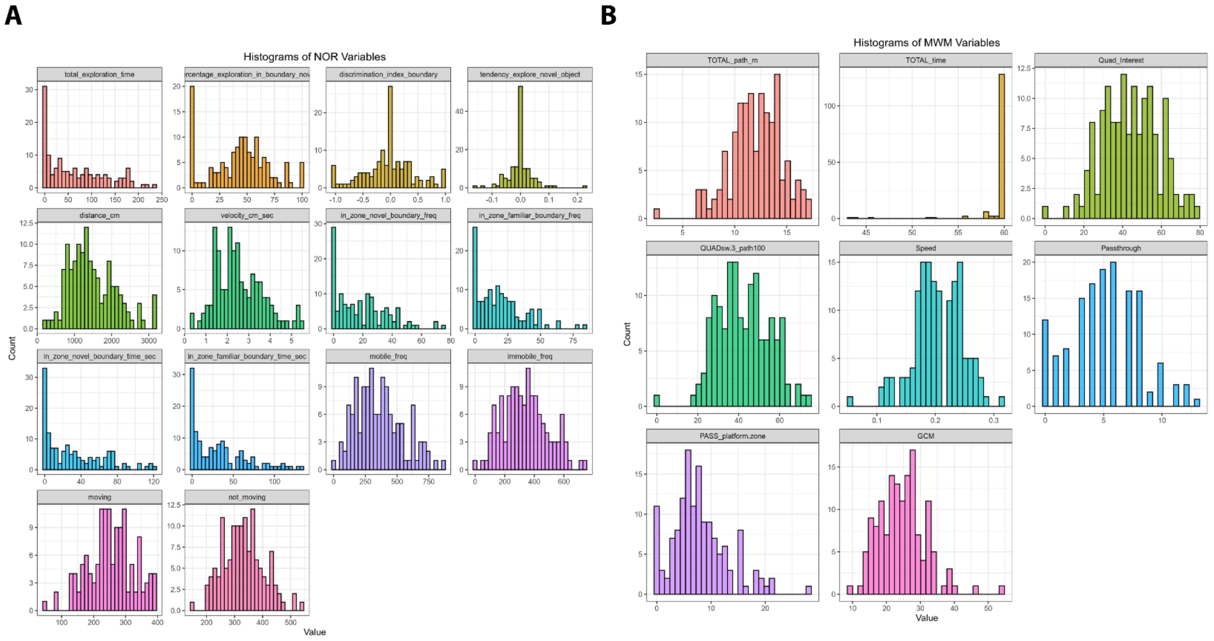


**Fig. S7. Distribution of Behavioral Outcomes**. Histograms showing the empirical distribution for all variables measured in the (A) Novel Object Recognition (NOR) and (B) Morris Water Maze (MWM) tasks. Data were visually inspected and tested for normality (Shapiro-Wilk) and homogeneity of variance (Levene's test) to inform model selection (e.g., Beta, GLM, or Hurdle models). See Tables S4-3 for details on selected model families.

**Table S1. Model selection per outcome in NOR.**

| **Variable** | **Variable type & distribution** | **Levene's Test (Homogeneity of variance)** | **Shapiro's test (Normality of residuals)** | **Selected model (distribution, link)** | **Significant term(s)** | **Interpretation** | **Final Model** |
|---|---|---|---|---|---|---|---|
| total_exploration_time | Continuous, right-skewed | Passed | Fail | Hurdle model (logistic + Gamma log) | WT Male in Exercise intervention | Among WT male mice, assignment to the Exercise intervention is associated with a model-predicted decrease of 41.7 seconds in the total exploration time relative to Control (95% CI [-76.2, -7.3], p = 0.02) | ~ Intervention * Genotype * Sex |
| percentage_exploration_in_boundary_novel | Proportion, spike at 0 | Not applicable | Not applicable | Hurdle model (logistic + Beta regression) | Not significant | | ~ Intervention * Genotype * Sex |
| discrimination_index_boundary | Continuous, bounded, spike at 0 | Passed | Fail | Hurdle model (logistic + Beta regression) | Not significant | | ~ Intervention * Genotype * Sex |
| tendency_explore_novel_object | Continuous, bounded, spike at 0 | Passed | Fail | Hurdle model (logistic + Beta regression) | Not significant | | ~ Intervention * Genotype * Sex |

| | | | | | | | |
|---|---|---|---|---|---|---|---|
| distance_cm | Continuous, bell-shaped | Passed | Fail | Gaussian, Identity | WT Female in combined intervention (Both) | Among WT female mice, assignment to the combined intervention is associated with a model-predicted decrease of 950.24 cm in the distance traveled relative to the Control intervention (95% CI [-1662.2, -238.3], p = 0.009). | ~ Intervention * Genotype * Sex |
| velocity_cm_sec | Continuous, bell-shaped | Passed | Passed | Linear Model | WT Male in Exercise WT Female in Both | Assignment to the combined intervention is associated with a model-predicted decrease of 1.58 cm/s in the distance traveled relative to Control (95% CI [-2.78, -0.37], p = 0.01). Assignment to the exercise intervention is associated with a model-predicted decrease of 1.18 cm/s in the distance traveled relative to Control in WT Male mice (95% CI [-2.29, -0.05], p = 0.04). | ~ Intervention * Genotype * Sex |
| In_zone_novel_boundary_freq | Count, spike at 0 | Irrelevant | Irrelevant | Hurdle model (logistic + Poisson / NegBin) | isolated significant term: GenotypeTg | Across all interventions and sexes, 3xTgAD animals differ (p = 0.03) | ~ Intervention * Genotype * Sex |
| In_zone_familiar_boundary_freq | Count, spike at 0 | Irrelevant | Irrelevant | Hurdle model (logistic + Poisson / NegBin) | male WT mice in Exercise | Assignment to the exercise intervention is associated with a model-predicted decrease of 14.8 counts in the in_zone_familiar_boundary_freq relative to Control (95% CI [-27.68, -1.92], p = 0.02). | ~ Intervention * Genotype * Sex |
| In_zone_novel_boundary_time_sec | Continuous, spike at 0 | Passed | Fail | Hurdle model (logistic + Gamma log) | Not significant | | ~ Intervention * Genotype * Sex |
| In_zone_familiar_boundary_time_sec | Continuous, right-skewed | Passed | Fail | Hurdle model (logistic + Gamma log) | male WT mice in Exercise | Assignment to the exercise intervention is associated with a model-predicted decrease of -24.38 seconds in the in_zone_familiar_boundary_time_sec relative to Control (95% CI [-45.96, -2.81], p = 0.03). | ~ Intervention * Genotype * Sex |
| mobile_freq | Count, bell-shaped | Irrelevant | Irrelevant | Negative Binomial | Not significant | | ~ Intervention * Genotype * Sex |
| immobile_freq | Count, bell-shaped | Irrelevant | Irrelevant | Negative Binomial | Both Male and Female in Tg mice in the HighFat Diet group | Assignment to the HighFat diet group is associated with a model-predicted increase of 149.56 counts in the immobilized frequency relative to Control in 3xTgAD female mice (95% CI [ 9.63, 289.48], p = 0.04), and an increase of 141.57 counts in 3xTgAD male mice (95% CI [13.13, 270.02], p = 0.03). | ~ Intervention * Genotype + Sex |
| moving | Continuous, bell-shaped | Passed | Passed | Linear Model | WT female mice in the combined intervention | Assignment to the combined intervention is associated with a model-predicted decrease of -96.79 seconds in the moving time relative to Control in WT female mice (95% CI [-179.15, -14.43], p = 0.02). | ~ Intervention * Genotype * Sex |
| not_moving | Continuous, bell-shaped | Passed | Passed | Linear Model | WT Male in Exercise intervention and WT Female in Combined intervention | Assignment to the exercise intervention is associated with a model-predicted increase of 96.75 seconds in the not-moving time relative to Control in WT male mice (95% CI [20.76, 172.74], p = 0.01). Assignment to the combined intervention is associated with a model-predicted increase of 94.05 seconds in the not-moving time relative to Control in WT female mice (95% CI [11.98, 176.13], p = 0.02). | ~ Intervention * Genotype * Sex |

The table provides a summary of model selection and significant effects for behavioral outcomes from the Novel Object Recognition (NOR) test.

**Table S2. Model selection per outcome in MWM.**

| Variable | Variable type & distribution | Levene's Test | Shapiro's test | Selected model (distribution, link) | Significant term(s) | Interpretation | Final Model |
|---|---|---|---|---|---|---|---|
| TOTAL_path_m | Continuous. Left-skewed. Total path swam in meters. | Passed | Failed | Log-transformed Gaussian | Weight | Body weight was included as a covariate to control for individual differences in locomotor capacity, but it is not a main outcome of interest. | Intervention * Genotype + Sex + Software + Weight |
| Quad_Interest | Continuous proportion (0–100%), mild Left-skewed. Percentage of time spent in the quadrant of interest (southwest, where the platform was located during training). | Not applicable. Levene's and Shapiro tests are Gaussian-specific. | Not applicable. Levene's and Shapiro tests are Gaussian-specific. | Beta regression | Several | WT male, Combined: +19.5% (0.194, 95% CI 0.052–0.34, p=0.007) WT male, HighFat: -17.7% (-0.177, 95% CI -0.31–-0.04, p=0.009) Tg female, Combined: -20.0% (-0.20, 95% CI -0.36–-0.03, p=0.02) Tg female, Diet: -14.0% (-0.14, 95% CI -0.27–-0.01, p=0.03) Tg male, Combined: +19.0% (0.19, 95% CI 0.002–0.39, p=0.047) | Intervention * Genotype * Sex + Software + Weight |
| Speed | Continuous. Skewed. Average swimming speed (m/s) | Passed | Failed | Log-transformed Gaussian | Not significant | | |
| Passthrough | Count. Right-skewed. Count of passages above the platform location during training. | Not applicable. Levene's and Shapiro tests are Gaussian-specific. | Not applicable. Levene's and Shapiro tests are Gaussian-specific. | quasipoisson GLM due to slight overdispersion | Several | WT Female, Exercise vs Control: model-predicted increase of 4.33 (95% CI: 0.17 to 8.49, p = 0.04); WT Male, Combined vs Control: increase of 5.15 (95% CI: 2.29 to 8.02, p < 0.001); Tg Female, Diet vs Control: decrease of 3.04 (95% CI: -5.07 to -1.00, p = 0.003); Tg Male, Combined vs Control: increase of 5.59 (95% CI: 1.12 to 10.06, p < 0.01) | Intervention * Genotype * Sex + Software + Weight |
| PASS_platform.zone | Count. Right-skewed. Count of passages above the quadrant | Not applicable. Levene's and Shapiro tests are | Not applicable. Levene's and Shapiro tests are Gaussian-speci | Poisson GLM or Negative binomial if overdispersed | Several | WT Male, Combined vs Control: increase of 8.53 (95% CI: 3.72 to 13.34, p < 0.001); WT Male, Diet vs Control: increase | Intervention * Genotype * Sex + Software + Weight |

| | | | | | | | |
|---|---|---|---|---|---|---|---|
| | where the platform was located during training (southwest). | Gaussian-spe cific. | fic. | | | of 3.46 (95% CI: 0.008 to 6.91, p = 0.049); WT Male, Exercise vs Control: increase of 3.85 (95% CI: 0.36 to 7.35, p = 0.03); Tg Female, Combined vs Control: decrease of 5.06 (95% CI: -9.27 to -0.86, p = 0.02); Tg Female, Diet vs Control: decrease of 5.92 (95% CI: -9.18 to -2.65, p < 0.001); Tg Male, Combined vs Control: increase of 9.01 (95% CI: 0.91 to 17.11, p = 0.03). | |
| GCM | Continuous. Right-skewed. Gallagher's cumulative measure (meters x seconds) is the time-weighted average distance from the platform across the trial. | Passed | Failed | Log-transformed Gaussian | Several | WT Female, Diet vs Control: decrease of 0.3 (95% CI: -0.56 to -0.04, p = 0.02); WT Male, Combined vs Control: decrease of 0.4 (95% CI: -0.66 to -0.13, p = 0.003); Tg Female, Diet vs Control: increase of 0.23 (95% CI: 0.0012 to 0.46, p = 0.048) | Intervention * Genotype * Sex + Software + Weight |

The table provides a summary of model selection and significant effects for behavioral outcomes from the Morris Water Maze (MWM) test.

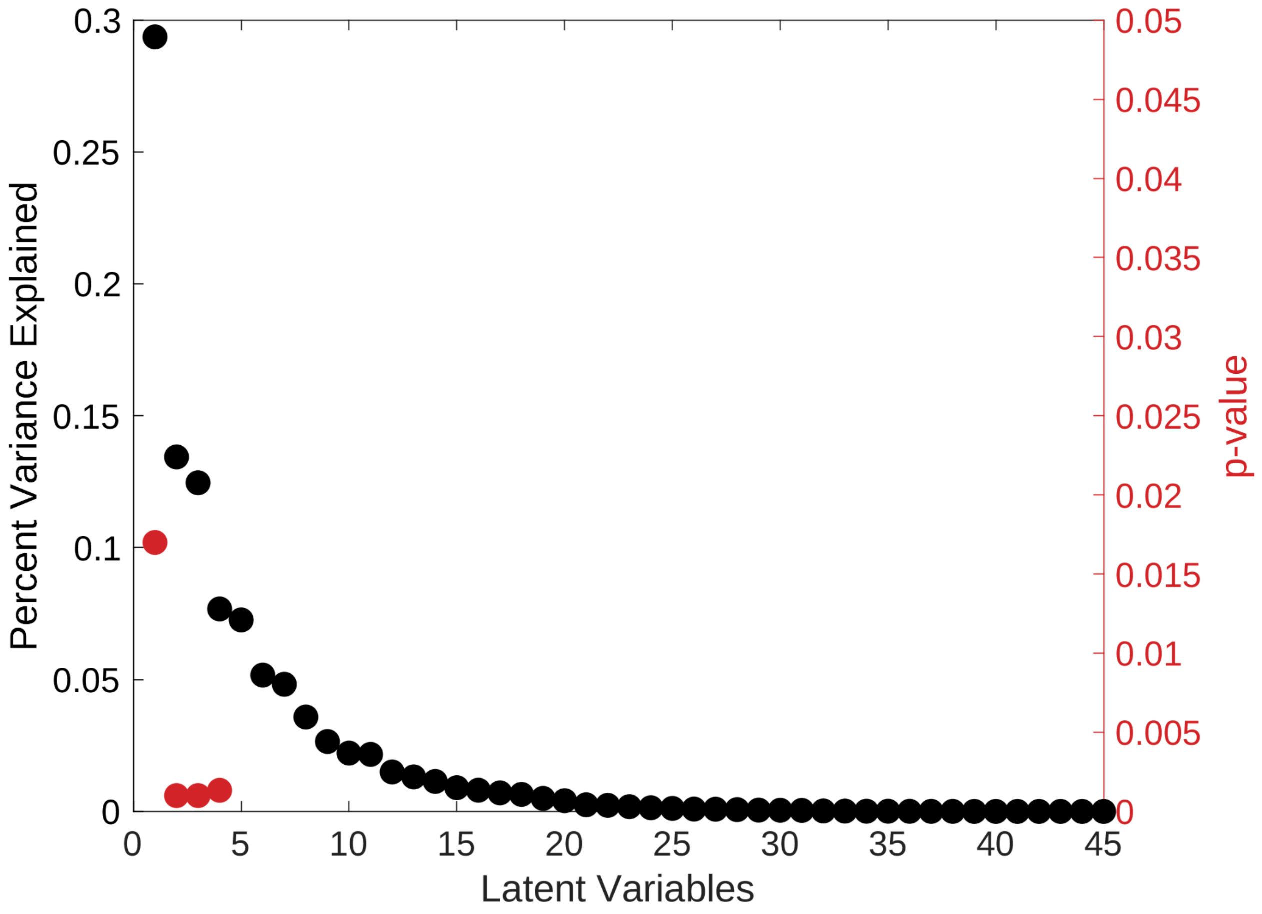


**Fig. S8. Significance and percentage of covariance explained by each latent variable.** PLS analysis resulted in 45 latent variables (LV). The graphic shows the percentage of variance explained (y-axis, left) and significant p-value (y-axis, right) per latent variable (x-axis). We obtained 4 significant latent variables that explained 29.36%, 13.44%, 12.46%, and 7.68% of the covariance, respectively.

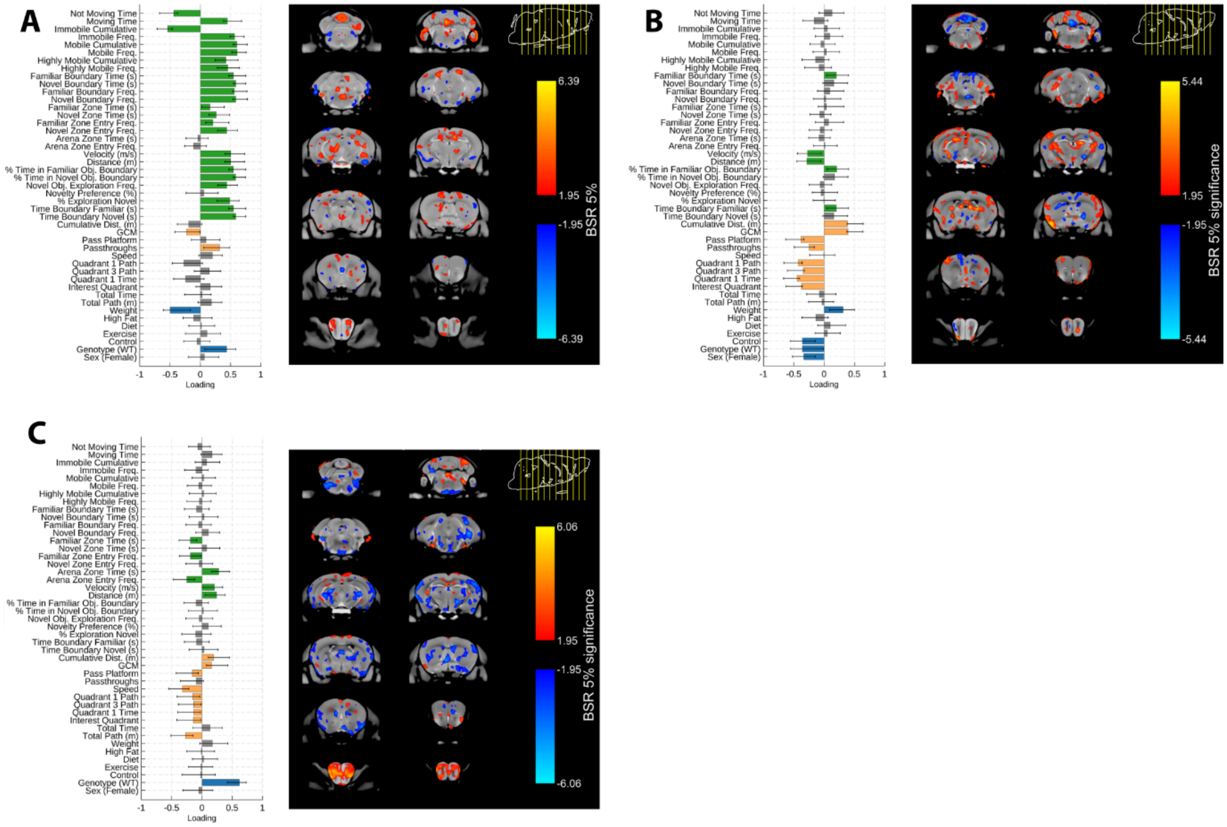


**Fig. S9. Behavioral weights and brain loadings for LV1, LV2, and LV4**. (A) Latent variable 1, (B) Latent variable 2, (C) Latent variable 4. To the left is the behavioral pattern that maximally covaries with the brain pattern. For the behavioral data, variables from the Novel Object Recognition test (NOR; in green), the Morris Water Maze (MWM; in yellow), and the treatment groups or demographics (in blue) are shown. The bars show how much each behavior contributes to the pattern of each LV (x-axis: correlation values). Singular value decomposition estimated the size of the bars, and confidence intervals were estimated by bootstrapping. Bars with error bars that cross the zero line are not significant (in gray). To the right, brain-loading bootstrap ratios (BSR) are overlaid on the population average. Positive BSR in orange indicates larger volumes, and negative BSR in blue indicates smaller volumes. BSR thresholded to a 95% confidence interval. N=96 mice with complete data.

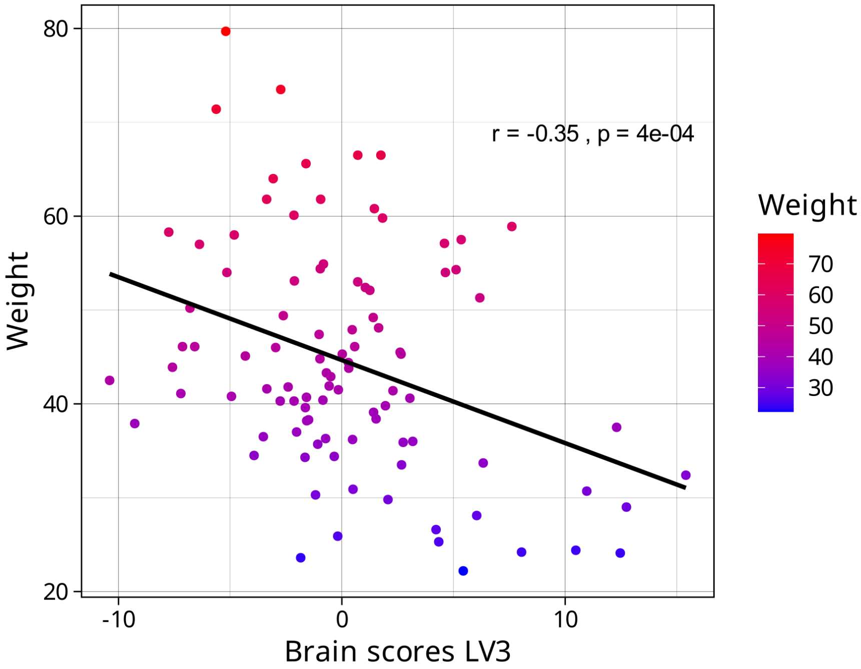


**Fig. S10. Relationship between Brain scores for LV3 and weight.** Brain scores (x-axis) reflect how much each mouse expresses the brain pattern observed in LV3. The negative relationship between weight in grams (y-axis) and brain scores shows that mice with higher weight are less represented (lower brain scores) by the brain pattern of LV3. Pearson's r = -0.35, p<0.5.

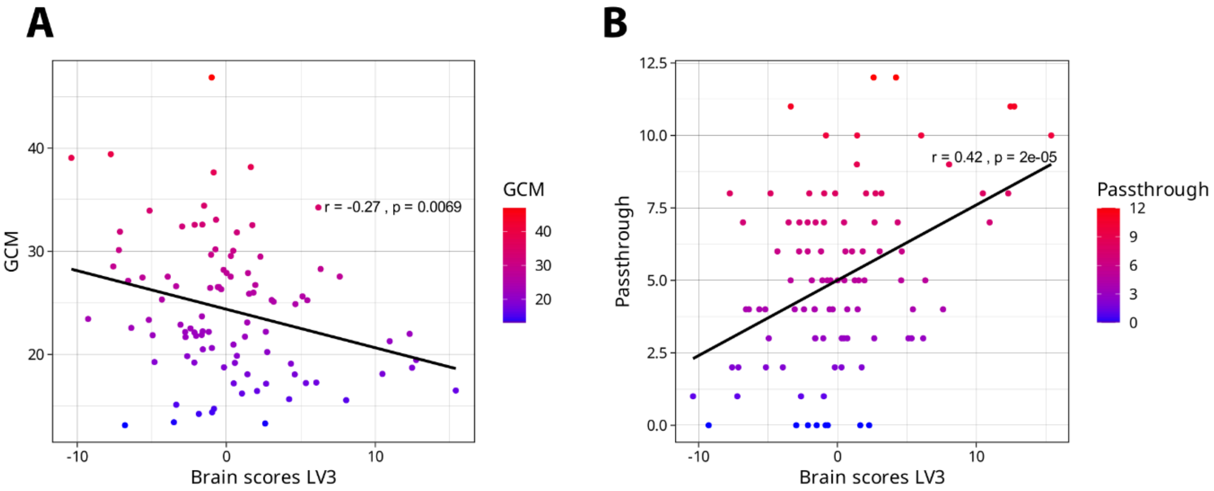


**Fig. S11. Relationship between Brain scores for LV3 and GCM, and Passthrough.** The scatter plot shows the relationship between Brain scores LV3 and GCM (A) and Passthrough (B). Left: the points are colored on a gradient, with lower values of GCM in blue and higher values in red. The correlation coefficient (r = -0.27) indicates a negative linear relationship between the variables, meaning that as the brain scores for LV3 increase, the GCM tends to be smaller. A smaller GCM indicates better memory performance. Right: the scatter plot shows the relationship between Brain scores LV3 and passthrough. The points are colored on a gradient, with lower Passthrough values in blue and higher values in red. The correlation coefficient (r = 0.42) indicates a positive linear relationship between the two variables, meaning that as brain scores increase, the passthrough also tends to increase. Higher pass-through counts suggest that the animal has accurately learned and remembers the platform's location. GCM: Gallagher's cumulative measure.

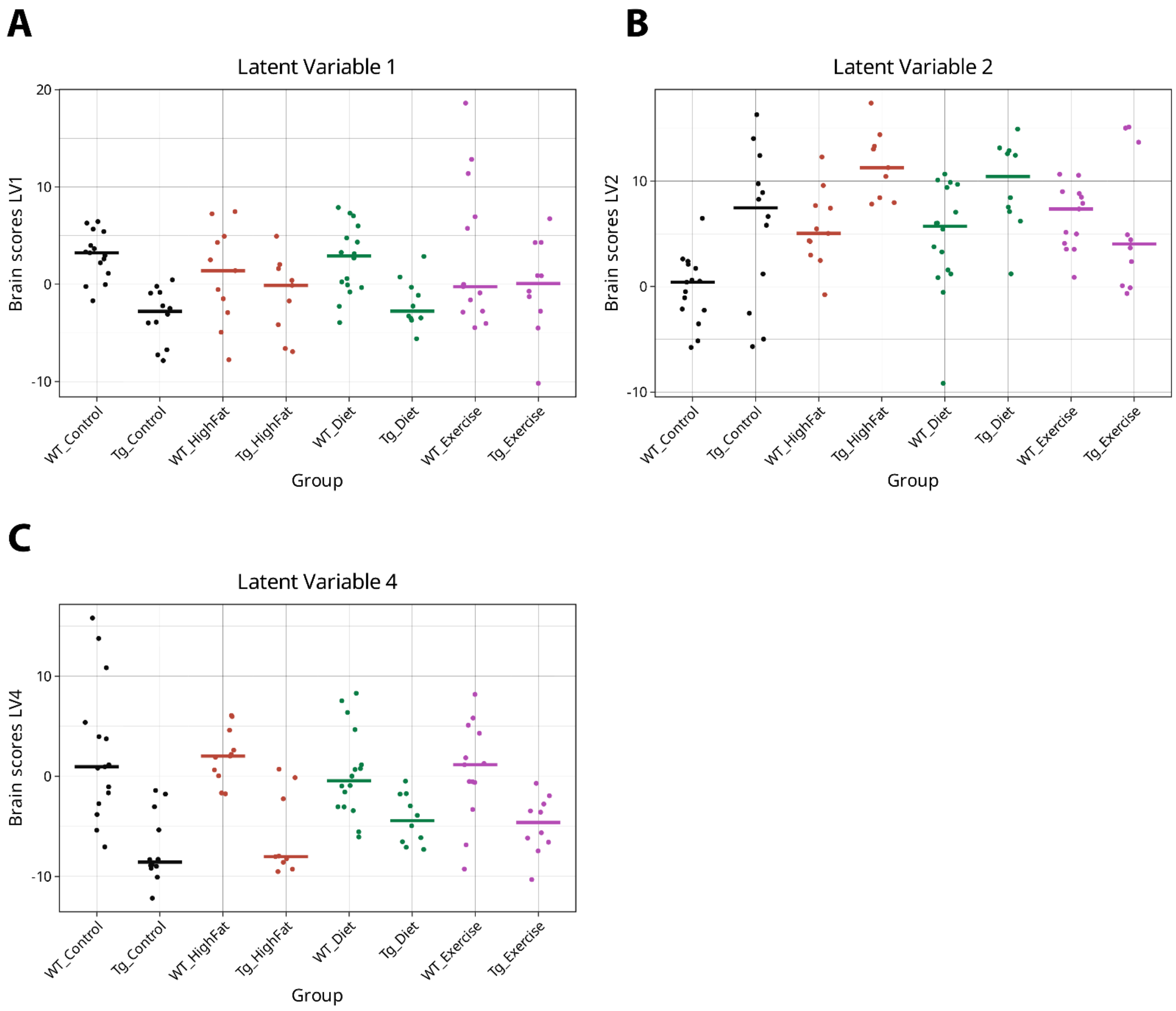


**Fig. S12. Brain scores and medians per group for LV1, 2, and 4.** Brain scores (x-axis) reflect how much each mouse expresses the brain pattern observed for LV1 (A), LV2 (B), and LV3 (C).

**Table S3. Nutrient information per diet.**

| Diet | Standard diet | Low-fat diet | High-fat diet |
|---|---|---|---|
| Energetic value (kcal/g) | 3.0 | 3.6 | 5.1 |
| Protein intake (%) | 16.4% | 20.50% | 18.30% |
| Protein source | - | casein | casein |
| Fat intake (%) | 3.7% | 10.50% | 60.30% |
| Fat source | Soybean oil | soybean oil and lard | soybean oil and lard |
| Carbohydrate intake (%) | 48.5% | 69.10% | 21.40% |
| Carbohydrate source | - | corn starch | maltodextrin |

From weaning until two months of age, mice were fed a standard diet (2016 Teklad Global 16% Protein Rodent Diet, Inotiv, Madison, Wisconsin USA; link), containing 4% fat from soybean oil and formulated for growth and maintenance. From two to four months of age, 80% of both 3xTgAD and WT mice were randomly assigned to an obesity-inducing high-fat diet (Teklad high-fat diet TD.06414 with 60% of kcal from fat, primarily lard, Inotiv, Madison, Wisconsin USA; link), while the remaining mice continued on an ingredient-matched low-fat diet (Teklad Purified Control Diet TD.08806 with 10.5% of kcal from fat, primarily soybean oil and lard, Inotiv, Madison, Wisconsin USA; link).

**Table S4. Number of mice per time point.**

| Group | Genotype | Intervention | Diet 0-2 mo. | Diet 2-4 mo. | Diet 4-6 mo. | TP 1 N (males, females) | TP 2 N (males, females) | TP 3 N (males, females) |
|---|---|---|---|---|---|---|---|---|
| **WT_Control** | WT | None | Standard | Low-fat | Low-fat | 16 (7, 9) | 15 (6, 9) | 15 (6, 9) |
| **WT_HighFat** | WT | None | Standard | High-fat | High-fat | 15 (9, 6) | 15 (9, 6) | 15 (9, 6) |
| **WT_Diet** | WT | Diet | Standard | High-fat | Low-fat | 17 (10, 7) | 17 (10, 7) | 17 (10, 7) |
| **WT_Exercise** | WT | Exercise | Standard | High-fat | High-fat | 18 (12, 6) | 18 (12, 6) | 18 (12, 6) |
| **WT_Both** | WT | Both | Standard | High-fat | Low-fat | 17 (10, 7) | 17 (10, 7) | 17 (10, 7) |
| **Tg_Control** | 3xTgAD | None | Standard | Low-fat | Low-fat | 16 (6, 10) | 16 (6, 10) | 16 (6, 10) |
| **Tg_HighFat** | 3xTgAD | None | Standard | High-fat | High-fat | 16 (10, 6) | 16 (10, 6) | 16 (10, 6) |
| **Tg_Diet** | 3xTgAD | Diet | Standard | High-fat | Low-fat | 16 (6, 10) | 16 (6, 10) | 16 (6, 10) |
| **Tg_Exercise** | 3xTgAD | Exercise | Standard | High-fat | High-fat | 18 (11, 7) | 17 (10, 7) | 16 (9, 7) |
| **Tg_Both** | 3xTgAD | Both | Standard | High-fat | Low-fat | 16 (9, 7) | 13 (6, 7) | 10 (5, 5) |
| | | | | | Total = | 165 | 160 | 156 |

The table lists the different diets each group received during the experiment and the number of mice available at each time point (TP 1: 2 months old, TP 2: 4 months old, and TP 3: 6 months old) when brain images were acquired. Out of the initial 165 mice involved (around 16 mice per group and a nearly equal number of males and females), 9 died or were sacrificed before the final neuroimaging timepoint, primarily due to fatal injuries as a result of fighting (four males and one female) which was precipitated primarily by the introduction of the running wheel (fighting caused an entire cage of 3xTgAD mice in the combined intervention to be sacrificed), but also from hydrocephalus (one male, one female), and complications as a result of MnCl2 injections (which were needed for improved MRI contrast) (two males).

**Table S5. List of regions (labels) from the olfactory bulb removed from the volumetric analysis.**

| Structure | Right label | Left label |
|---|---|---|
| Olfactory bulb: glomerular layer | 337 | 345 |
| Olfactory bulb: external plexiform layer | 338 | 346 |
| Olfactory bulb: mitral cell layer | 339 | 347 |
| Olfactory bulb: internal plexiform layer | 340 | 348 |
| Olfactory bulb: granule cell layer | 341 | 349 |
| Accessory olfactory bulb: glomerular, external plexiform and mitral cell layer | 342 | 350 |
| Accessory olfactory bulb: granule cell layer | 343 | 351 |
| Anterior olfactory nucleus | 344 | 352 |
| olfactory peduncle | 5 | 105 |
| olfactory tubercle | 95 | 145 |

By removing olfactory bulb labels, the number of scans that passed QC increased, and this was preferable to retain a larger N per group.

**Table S6. Number of mice per group for NOR and MWM.**

| Group | Genotype | Intervention | Diet 0-2 mo. | Diet 2-4 mo. | Diet 4-6 mo. | MWM N (males, females) | NOR N (males, females) |
|---|---|---|---|---|---|---|---|
| **WT_Control** | WT | None | Standard | Low-fat | Low-fat | 15 (9, 6) | 15 (9, 6) |
| **WT_HighFat** | WT | None | Standard | High-fat | High-fat | 14 (6, 8) | 14 (5, 9) |
| **WT_Diet** | WT | Diet | Standard | High-fat | Low-fat | 17 (7, 10) | 17 (7, 10) |
| **WT_Exercise** | WT | Exercise | Standard | High-fat | High-fat | 18 (6, 12) | 13 (6, 7) |
| **WT_Both** | WT | Both | Standard | High-fat | Low-fat | 17 (7, 10) | 8 (4, 4) |
| **Tg_Control** | 3xTgAD | None | Standard | Low-fat | Low-fat | 14 (9, 5) | 16 (10, 6) |
| **Tg_HighFat** | 3xTgAD | None | Standard | High-fat | High-fat | 13 (3, 10) | 14 (4, 10) |
| **Tg_Diet** | 3xTgAD | Diet | Standard | High-fat | Low-fat | 16 (10, 6) | 14 (10, 4) |
| **Tg_Exercise** | 3xTgAD | Exercise | Standard | High-fat | High-fat | 13 (7, 6) | 15 (6, 9) |
| **Tg_Both** | 3xTgAD | Both | Standard | High-fat | Low-fat | 8 (4, 4) | 7 (3, 4) |
| | | | | | Total = | 145 | 133 |

The table provides the number of mice per sex included in the Morris Water Maze (MWM) and Novel Object Recognition (NOR) statistical analyses (mice with complete data). The behavioural tests were conducted at 6 months of age, following the last MRI scan.
